# Compositional heterogeneity and outgroup choice influence the internal phylogeny of the ants

**DOI:** 10.1101/173393

**Authors:** Marek L. Borowiec, Christian Rabeling, Seán G. Brady, Brian L. Fisher, Ted R. Schultz, Philip S. Ward

## Abstract

Knowledge of the internal phylogeny and evolutionary history of ants (Formicidae), the world’s most species-rich clade of eusocial organisms, has dramatically improved since the advent of molecular phylogenetics. A number of relationships at the subfamily level, however, remain uncertain. Key unresolved issues include placement of the root of the ant tree of life and the relationships among the so-called poneroid subfamilies. Here we assemble a new data set to attempt a resolution of these two problems and carry out divergence dating, focusing on the age of the root node of crown Formicidae. For the phylogenetic analyses we included data from 110 ant species, including the key species *Martialis heureka*. We focused taxon sampling on non-formicoid lineages of ants to gain insight about deep nodes in the ant phylogeny. For divergence dating we retained a subset of 62 extant taxa and 42 fossils in order to approximate diversified sampling in the context of the fossilized birth-death process. We sequenced 11 nuclear gene fragments for a total of ~7.5 kb and investigated the DNA sequence data for the presence of among-taxon compositional heterogeneity, a property known to mislead phylogenetic inference, and for its potential to affect the rooting of the ant phylogeny. We found sequences of the Leptanillinae and several outgroup taxa to be rich in adenine and thymine (51% average AT content) compared to the remaining ants (45% average). To investigate whether this heterogeneity could bias phylogenetic inference we performed outgroup removal experiments, analysis of compositionally homogeneous sites, and a simulation study. We found that compositional heterogeneity indeed appears to affect the placement of the root of the ant tree but has limited impact on more recent nodes. We put forward a novel hypothesis regarding the rooting of the ant phylogeny, in which *Martialis* and the Leptanillinae together constitute a clade that is sister to all other ants. After correcting for compositional heterogeneity this emerges as the best-supported hypothesis of relationships at deep nodes in the ant tree. The results of our divergence dating under the fossilized birth-death process and diversified sampling suggest that the crown Formicidae originated during the Albian or Aptian ages of the Lower Cretaceous (103–124 Ma). In addition, we found support for monophyletic poneroids comprising the subfamilies Agroecomyrmecinae, Amblyoponinae, Apomyrminae, Paraponerinae, Ponerinae, and Proceratiinae, and well-supported relationships among these subfamilies except for the placement of Proceratiinae and (Amblyoponinae + Apomyrminae). Our phylogeny also highlights the non-monophyly of several ant genera, including *Protanilla* and *Leptanilla* in the Leptanillinae, *Proceratium* in the Proceratiinae, and *Cryptopone*, *Euponera*, and *Mesoponera* within the Ponerinae.

## 1. Introduction

Ants are among the world’s dominant social insects, with more species and greater ecological impact than any other group of eusocial animals (Hölldobler and Wilson, 1990). Knowledge of ant phylogeny is vital to understanding the processes driving the evolution of these ubiquitous and diverse organisms.

About 90% of extant ant diversity, or almost 11,000 species and 9 of 16 subfamilies, belongs to a group dubbed the ”formicoid clade” (Brady et al., 2006). The relationships among the subfamilies of the formicoid clade are well-resolved. This contrasts with the uncertain branching order of lineages outside of formicoids. Two enduring issues in higher ant phylogeny, highlighted by recent studies (Moreau et al., 2006; Brady et al., 2006; Rabeling et al., 2008) and a review by Ward (2014), include the identity and composition of the lineage that is sister to all other extant ants, and whether the so-called poneroid subfamilies (Agroecomyrmecinae, Amblyoponinae, Apomyrminae, Paraponerinae, Ponerinae, Proceratiinae) form a clade or a grade.

In molecular phylogenetic studies published to date, two ant subfamilies, the Martialinae and Leptanillinae (recently redefined to include the genus *Opamyrma*; Ward and Fisher (2016)), have been competing for the designation as the sister group to all other ants (Figure 1) (Rabeling et al., 2008; Kück et al., 2011; Moreau and Bell, 2013; Ward and Fisher, 2016). Rabeling et al. (2008) discovered and described *Martialis heureka*, and inferred that this sole species of the morphologically divergent Martialinae is the sister species to all other ants. Subsequent phylogenies, however, proposed that either the subfamily Leptanillinae is the lineage first branching from the rest of the ants (Kück et al., 2011), or were ambiguous about the placement of the ant root (Moreau and Bell, 2013).

**Figure 1:**
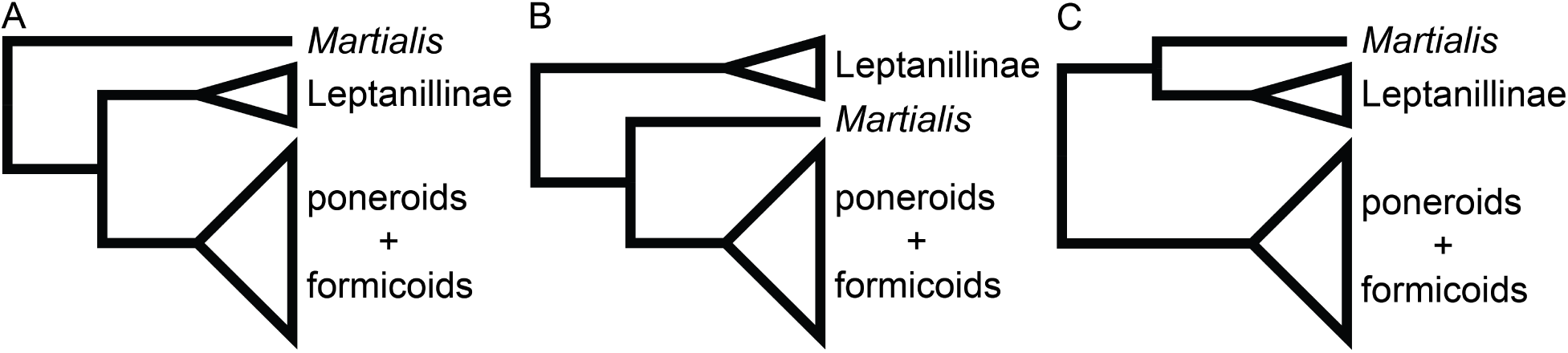
Comparison of alternative hypotheses for the root of the ant phylogeny. A, *Martialis* sister to all other ants (Rabeling et al., 2008); B, Leptanillinae sister to all other ants, including *Martialis* (Kück et al., 2011); C, *Martialis* plus Leptanillinae sister to all other ants (this study).

The monophyly of the poneroid subfamilies was recovered by Moreau et al. (2006) but it was later contested by Brady et al. (2006) and Rabeling et al. (2008). Brady et al. (2006) pointed out that long-branch attraction may be responsible for poneroid monophyly and found that non-monophyly of poneroids could not be rejected based on their data set of seven nuclear gene fragments. Most subsequent phylogenetic studies failed to satisfactorily resolve this issue (reviewed in Ward (2014)) although poneroid monophyly was recovered with strong support by Ward and Fisher (2016). A recently published phylogenomic study also supported poneroid monophyly (Branstetter et al., 2017b). The phylogenomic data were apparently insufficient, however, to resolve relationships among poneroid lineages because of low taxon sampling (Branstetter et al., 2017b).

In addition to the phylogenetic uncertainty present in the above mentioned studies, there is also potential for systematic bias to preclude correct inference of ant relationships, especially near the root of the ant tree (Ward, 2014). This is because most ants are relatively guanine and cytosine-(GC) rich compared to many aculeate outgroups and species of the Leptanillinae, which are unusually adenine and thymine-(AT) rich. Such compositional heterogeneity is known to mislead phylogenetic inference (Jermiin et al., 2004) and most of the commonly used models of sequence evolution do not take it into account. This and other potential violations may lead to poor model fit, which was demonstrated in phylogenetics in general (Brown, 2014) and in an ant phylogeny specifically (Rabeling et al., 2008). It is thus possible that the basal position of the Leptanillinae appearing in some studies is an artifact resulting from model misspecification.

To address these issues of uncertainty and potential bias near the base of the ant tree of life, we assembled a new comprehensive data set that included all ant subfamilies, with sampling focused on non-formicoid lineages. The amount of sequence data we gathered is significantly greater than in previous studies that included the pivotal species *Martialis heureka*. We investigated the potential of these data to be biased by base-frequency heterogeneity and implemented strategies aimed at minimizing such bias.

Another outstanding question concerns the age of the most-recent common ancestor of ants, which has been variously estimated to have lived as recently as ca. 115 Ma (Brady et al., 2006) or as early as 168 Ma (Moreau et al., 2006), with the oldest undisputed crown formicid fossil *Kyromyrma neffi* dated at 92 Ma (Grimaldi and Agosti, 2000; Barden, 2017). Efforts to infer the age of origin of crown ants have been conducted thus far using either penalized likelihood (Sanderson, 2002) or node dating in a Bayesian framework (Drummond and Rambaut, 2007). Here we take advantage of the fossilized birth-death process framework (Heath et al., 2014), which was recently extended to accommodate diversified taxon sampling (Zhang et al., 2016).

The recently developed fossilized birth-death process (FBD) approach to divergence dating provides several advantages over the node-dating approach, as it explicitly treats both extant and fossil taxa as parts of the same underlying diversification process. This is different from the node-dating approach, where fossils only provide clues about the probability-density distributions of ages for certain splits (Heath et al., 2014). FBD is thus able to avoid some of the challenges documented for node-dating, such as formulation of the statistical problem accidentally precluding reasonable age estimate for a node of interest (Brown and Smith, 2017). Early implementations of the FBD, however, shared the assumption of other Bayesian divergence-time estimation methods that taxon sampling reflects complete or random sampling of the diversification process that created the phylogeny. Because phylogenies of higher taxa aim for maximizing phylogenetic diversity (Höhna et al., 2011), this assumption is often violated and is known to cause biased age estimates (Beaulieu et al., 2015). The most recent implementation of the FBD accounts for this by explicitly allowing modeling under a diversified sampling scheme (Zhang et al., 2016).

## 2. Methods

### 2.1. Taxon sampling and data collection

We collected data from 11 nuclear loci for 110 ant species, including *Martialis heureka*, and 13 outgroup taxa. Our data set has extensive sampling within Leptanillinae (21 species), all major lineages of the poneroids (66 species), and at least one representative of all formicoid subfamilies (22 species). The sequence data comes from *28S ribosomal DNA* (*28S*) and ten nuclear protein-coding genes: *abdominal*-*A* (*abdA*), *elongation factor 1*-*alpha F2 copy* (*EF1aF2*), *long wavelength rhodopsin* (*LW Rh*), *arginine kinase* (*argK*), *topoisomerase I* (*Top1*), *ultrabithorax* (*Ubx*), *DNA pol*-*delta* (*POLD1*), *NaK ATPase* (*NaK*), *antennapedia* (*Antp*), and *wingless* (*Wg*). Sequences were assembled with Sequencher v5.2.2 (Gene Codes Corporation, Ann Arbor, MI, U.S.A.), aligned with Clustal X v2.1 (Thompson et al., 1997), and manually edited and concatenated with MacClade v4.08 (Maddison and Maddison, 2005). We excluded introns, autapomorphic indels (9 bp *abdA*, 3 bp *EF1aF2*, 9 bp *Ubx*, 3 bp *POLD1*, 3 bp *NaK*, 75 bp *Antp*, 3 bp *Wg*), and hypervariable regions of *28S*. The resulting data matrix is 7,451 bp long and has no missing data except for gaps introduced by non-autapomorphic indels, which constitute 3% of the data. Our protocols for extraction and amplification were described in detail in Ward et al. (2010) and Ward and Fisher (2016). Of the 1,353 sequences, 676 were newly generated for this study (GenBank accessions XXXXXXXX-XXXXXXXX [submission in progress]; Supplementary Table 4).

### 2.2. Compositional heterogeneity

We assessed compositional heterogeneity using methods implemented in the p4 phylogenetics library (Foster, 2004). To investigate whether our data significantly departed from the assumption of homogeneity across taxa we applied two tests: 1) a *χ*^2^ test for compositional heterogeneity, and 2) a more sensitive, simulation-based test corrected for phylogeny. The phylogeny-corrected tests involved inferring a neighbor-joining tree for the data, constructed with BioNJ (Gascuel, 1997), onto which we simulated 1,000 replicates using GTR+4Γ model. The distribution of sequence compositions in the empirical data were then compared against simulated data. Following the finding that the combined data set failed both the *χ*^2^ and phylogenetically-corrected tests, we further divided the data set into partitions as follows: 1) each locus as its own partition, and 2) first, second, and third codon position within each locus assigned to separate partitions. We then ran the two compositional heterogeneity tests on each partition separately and recorded the results. We also computed base frequencies for each taxon and partition using AMAS (Borowiec, 2016).

### 2.3. Data matrices

In addition to the full data set, we composed two different matrices by excluding the most AT-rich and the most GC-rich outgroups, respectively. To this end, we first ranked non-ant taxa by AT content and then removed six taxa that had either the highest or lowest AT content, which also corresponded to above or below mean AT content for the entire data set. We retained moderately AT-rich *Pristocera* MG01 as the external outgroup.

We also constructed a data set from which we removed all heterogeneous partitions (i.e., *28S*, all third codon positions, and first codon positions of *POLD1* and *Ubx*).

Selected statistics of the four analyzed data matrices are in the first four rows of Table 1.

**Table 1:** Data properties.

| Data matrix | Length | Percent gaps | Variable sites | Parsimony informative sites |
| --- | --- | --- | --- | --- |
| Full data set | 7451 | 3.0 | 3902 | 3369 |
| AT-rich outgroups removed | 7451 | 3.0 | 3806 | 3237 |
| GC-rich outgroups removed | 7451 | 3.0 | 3825 | 3291 |
| Homogeneous partitions | 3995 | 2.7 | 1181 | 831 |
| 28S | 845 | 6.8 | 425 | 317 |
| abdA | 621 | 3.7 | 311 | 258 |
| argK | 673 | 0.3 | 334 | 296 |
| Antp | 865 | 14.0 | 497 | 397 |
| EF1aF2 | 517 | 0.0 | 206 | 194 |
| LW Rh | 458 | 0.0 | 287 | 262 |
| NaK | 955 | 0.0 | 409 | 376 |
| POLD1 | 583 | 0.0 | 374 | 325 |
| Top1 | 880 | 0.0 | 477 | 426 |
| Ubx | 630 | 0.7 | 313 | 280 |
| Wg | 424 | 3.4 | 269 | 238 |
| All 1st codon pos. | 2200 | 2.5 | 829 | 620 |
| All 2nd codon pos. | 2199 | 2.5 | 536 | 349 |
| All 3rd codon pos. | 2207 | 2.5 | 2112 | 2083 |

### 2.4. Partitioning

In the light of recent criticism of the k-means partitioning strategy (Baca et al., 2017), for each of the four data sets we used two different strategies for partitioning: a ’greedy’ algorithm on predefined user partitions and the k-means partitioning algorithm (Frandsen et al., 2015) as implemented in PartitionFinder 2 pre-release v13 (Lanfear et al., 2017). The greedy strategy relies on user-defined sets of characters as input and our predefined sets constituted sites from the three codon positions in each of the loci used, except for *28S* which was defined as a single set. We used a maximum-likelihood tree generated with the fast RAxML algorithm (”-f E” option) (Stamatakis, 2014) as the starting tree for each PartitionFinder run, which then used PhyML for subsequent steps of the algorithm (Guindon et al., 2010). Because of our use of MrBayes in downstream analyses, we restricted models to be evaluated by PartitionFinder to those available in that program.

### 2.5. Phylogenetics

We constructed phylogenies for all empirical data sets using Bayesian inference with MrBayes v3.2.6 (Ronquist et al., 2012) and maximum-likelihood with IQ-TREE v1.4.2 beta (Nguyen et al., 2014). For the Bayesian analyses, we ran two separate runs, four chains each, for 20 to 80 million generations for each of the eight analyses. We used a 20% burnin fraction and determined mixing and convergence by examining output of MrBayes ”sump” command, including average standard deviation of split frequencies (below < 0.01), effective sample size (ESS) for each parameter (minimum 200), and potential scale reduction factor (PSRF) near 1.0. In the maximum-likelihood analyses we specified the same partitioning schemes and substitution models as for the Bayesian inference. We changed the default IQ-TREE settings by using the slow nearest-neighbor interchange search (”-allnni” option) and setting the number of unsuccessful iterations to stop at 1,000 instead of the default 100 (”-numstop” option). We assessed support by running 2,000 ultrafast boostrap replicates (Minh et al., 2013). The authors of this fast bootstrap approximation point out that this algorithm tends to overestimate probability of a correct relationship under 70% support but is more unbiased than RAxML’s rapid bootstrapping above that threshold, resulting in 95% support being approximately equal to 95% probability of a relationship being true under the true model. Therefore, support at or above 95% should be interpreted as significant (Minh et al., 2013). We also attempted analyses under tree-heterogeneous models implemented in p4 (Foster, 2004) and nhPhyloBayes (Blanquart and Lartillot, 2008) on the full data matrix, but these turned out to be prohibitively computationally expensive, requiring by our estimate a minimum of five months to reach convergence.

### 2.6. Simulation

To further assess the sensitivity of the ant phylogeny to bias we simulated a data set with compositional heterogeneity comparable to that present in our data set. In particular, we were interested in investigating whether the position of *Martialis* could be incorrectly inferred as sister to the poneroids plus formicoids clade even if the data were simulated on a topology where *Martialis* is sister to Leptanillinae. To create the simulated data set we first split our empirical alignment, excluding ribosomal *28S*, into alignments of first, second, and third codon positions. We then used a fixed topology which had *Martialis* as sister to Leptanillinae, in this case the Bayesian posterior consensus from the full data set analysis under k-means partitioning, to estimate branch lengths for each of the three alignments. To estimate the branch lengths we used IQ-TREE (Nguyen et al., 2015) under the general time-reversible model with rate heterogeneity described by a proportion of invariant sites and a gamma distribution discretized into four bins (GTR+pinv+4Γ). For each of the codon position alignments we also calculated the proportion of invariant sites and base frequencies for all taxa using AMAS (Borowiec, 2016). Furthermore, we calculated average base frequencies for alignments composed of first and second positions. For the third codon positions alignment we calculated two average base frequencies: one for the 25 most AT-rich taxa, represented almost exclusively by Leptanillinae species and some of the outgroups, and the other for all remaining taxa, thus approximating mean base frequencies for AT-rich and GC-rich taxa, respectively. We then used the topologies with branch lengths, proportion of invariant sites, and empirical base frequencies to simulate three separate alignments, each 2,200 sites long, similar to the empirical data, under GTR+pinv+4Γ using p4 (Foster, 2004). For the alignments imitating first and second codon positions we simulated the data under a tree-homogeneous model, but for the third codon position alignment we used two composition vectors corresponding to the two empirical means of AT-rich (A: 0.24, C: 0.24, G: 0.23, T: 0.29) and GC-rich sequences (A: 0.16, C: 0.33, G: 0.31, T: 0.20) at that position. The AT-rich frequencies were applied to the Leptanillinae and outgroup taxa considered AT-rich in our outgroup taxa removal experiments outlined above. We replicated the simulation 100 times for each alignment under different starting seed numbers. We then performed maximum-likelihood analyses under GTR+pinv+4Γ (using IQ-TREE settings as described above) for the concatenated simulated data as well as each of the three simulated alignments separately. Following the inference we constructed majority-rule consensus trees using all 100 maximum-likelihood trees for all codon position simulations and the concatenated alignments in order to visualize the topology recovered from simulated alignments.

### 2.7. Divergence time estimation

We performed divergence dating under the fossilized birth-death process (Heath et al., 2014) and diversified sampling, as implemented in MrBayes v3.2.6 (Ronquist et al., 2012; Zhang et al., 2016). To obtain a taxon sample that most closely approximates diversified sampling *sensu* Höhna et al. (2011), i.e., one extant descendant sampled for each branch that was present at a given time in the past, we pruned our original alignment down to 62 species. To calibrate the analysis we used 42 fossil calibrations (See Supplementary Table 3) and a diffuse root node exponential prior with a mean of 250 Ma and offset at 150 Ma (just older than the oldest fossil calibration used). Because we did not include morphological data in our analysis to place the fossils (”total evidence” dating sensu Ronquist et al. (2012)), we assigned them to appropriate groups via monophyly constraints (Heath et al., 2014). For this analysis we also constrained the topology of outgroup taxa to correspond to the aculeate phylogeny recovered by a recent phylogenomic study (Branstetter et al., 2017a). We chose the topology from this study over that of Peters et al. (2017) because the latter did not include Rhopalosomatidae and Sierolomorphidae and thus arguably had insufficient taxon sampling to correctly place Vespidae.

We ran the analysis with four runs, each with six incrementally heated chains, for 100 million generations and discarded 10% of the samples as burn-in. The analysis was unpartitioned, with the GTR+6Γ substitution model and a relaxed independent clock model with rate drawn from a gamma distribution. We checked for convergence by examining the average standard deviation of split frequencies towards the end of the run (<0.006), potential scale reduction factor values for each parameter (maximum 1.028, average 1.001), and effective sample sizes (>500 for combined runs). We examined MCMC trace files with Tracer v1.5 to confirm that the two runs converged on all parameters and to compare posterior distributions to the analysis without data (i.e., under the prior).

### 2.8. Data availability

All data matrices, configurations for and output from PartitionFinder, Bayesian, and maximum-likelihood analyses, as well as custom scripts used are available the Zenodo data repository, DOI: 10.5281/zenodo.838799.

## 3. Results

We first summarize results regarding compositional heterogeneity, followed by a discussion of the placement of the formicid root as it was impacted by the different methods. Second, we present phylogenetic findings which were less sensitive to different analytical treatments, including poneroid monophyly, the relationships within poneroids, and non-monophyly of some of the non-formicoid genera.

### 3.1. Compositional heterogeneity

Within the ingroup, ants in the Leptanillinae stands out as particularly AT-rich at 50.9% on average compared to a mean of 46.2% for all taxa (or 44.9% for non-leptanilline ants). At 46.6%, *Martialis* is close to the mean (Figure 2, Supplementary Table 1).

**Figure 2:**
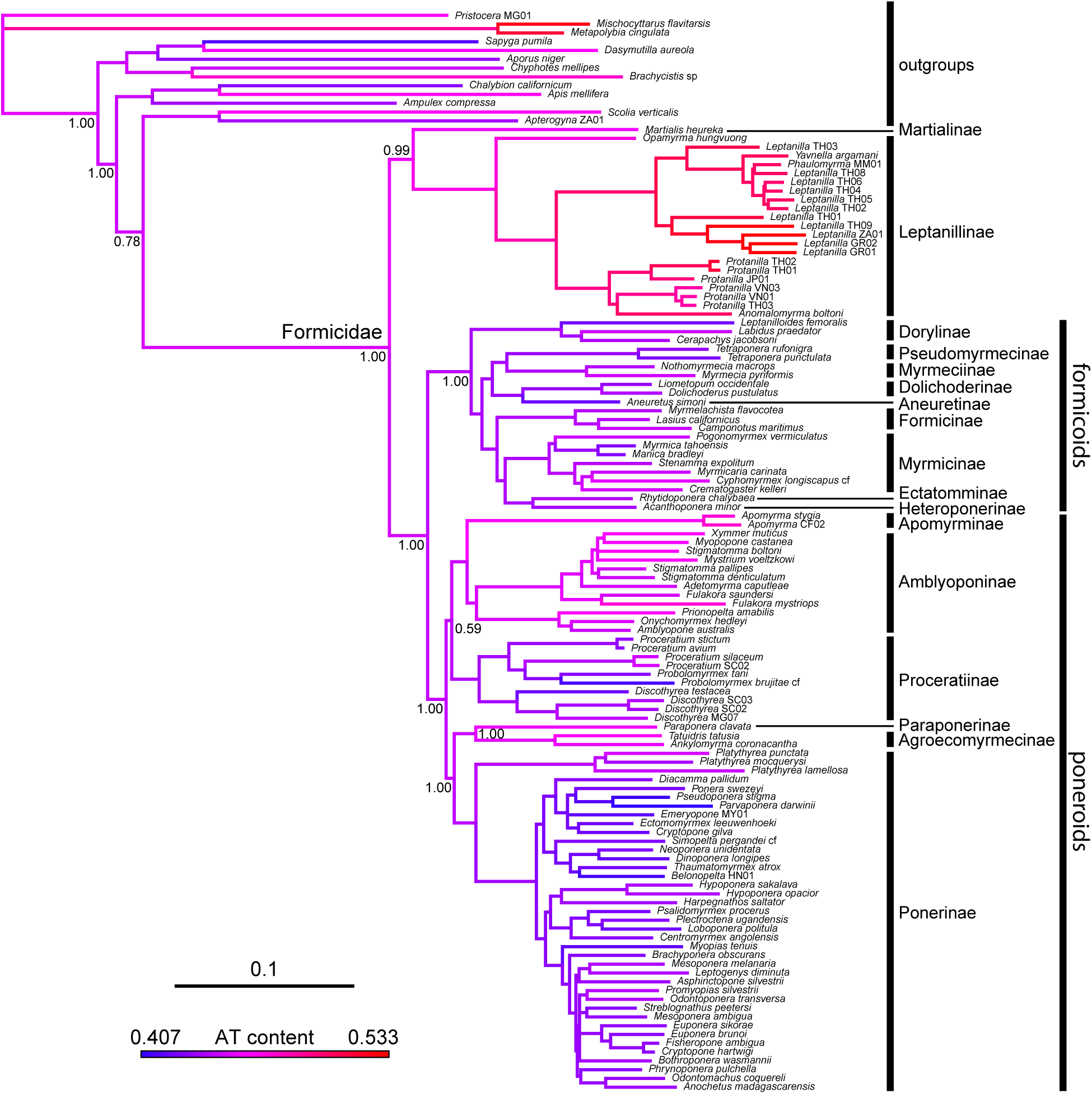
Preferred phylogenetic hypothesis for the ingroup (Formicidae), with AT content indicated for each terminal taxon. Tree topology with branch lengths from the Bayesian analysis of the full data matrix under k-means partitioning strategy. See Supplementary Figure 2 for support values on all nodes. Warmer branch colors signify higher AT content. Scale in expected substitutions per site.

We found considerable compositional heterogeneity among taxa in our data set, mostly confined to third codon positions. Overall difference in AT content among taxa was 12.6% across the entire data set. This difference was equal to 37.3% at third codon positions, compared to only 3.5% at first codon positions and 1.1% at the second codon positions. Third codon positions also accounted for 56.7% variable sites and 61.8% parsimony informative sites of the data set.

Consistent with this pattern, the phylogeny-corrected test identified all third codon partitions as those for which the homogeneity assumption could be rejected with a *p* < 0.05 (Supplementary Table 2). In addition to the third codon positions, *28S*, and first codon positions of *POLD1* and *Ubx* were found to violate the homogeneity assumption using this test. Similar to the phylogeny-corrected test, the *χ*^2^ test identified all of the third codon positions as heterogeneous but it did not reject the homogeneity assumption for *28S* and first codon positions of *POLD1* and *Ubx* (Supplementary Table 2).

Compositional heterogeneity was also high among the outgroups. Those that fell above the mean AT content for the entire alignment included (in order of decreasing AT content) *Mischocyttarus flavitarsis*, *Metapolybia cingulata*, *Brachycistis* sp., *Apis mellifera*, *Dasymutilla aureola*, *Scolia verticalis*, and *Pristocera* MG01. The outgroup species that were more GC-rich than average were (in order of decreasing AT content) *Chyphotes mellipes*, *Ampulex compressa*, *Apterogyna* ZA01, *Aporus niger*, *Chalybion californicum*, and *Sapyga pumila* (Supplementary Table 1).

### 3.2. Analyses of the full data matrix

With all taxa retained and no attempt at reducing the compositional heterogeneity of the data, the partitioning strategy has a strong effect on the results. Under the k-means strategy *Martialis heureka* is sister to the Leptanillinae with strong support of *pp* = 0.99 in the Bayesian analysis (Figure 2; Supplementary Figure 2) and low support of 87% bootstrap in the maximum-likelihood tree (Supplementary Figure 4). Under the greedy analyses, both Bayesian and maximum-likelihood trees recover a topology where the Leptanillinae are sister to the remaining Formicidae including *Martialis*, although support for this topology is below significance (*pp* = 0.81 and bootstrap 93%; Supplementary Figures 1 and 3).

### 3.3. Effects of outgroup removal

Relative to the full data matrix with all 13 outgroup species, removal of the AT-rich outgroups (*Apis mellifera*, *Brachycistis* sp., *Dasymutilla aureola*, *Scolia verticalis*, *Metapolybia cingulata*, and *Mischocyttarus flavitarsis*) shifts support towards a tree where *Martialis* and the Leptanillinae together form a clade that is sister to all other ants. In consensus Bayesian trees, the support for this clade is at *pp* = 0.91 under greedy (Supplementary Figure 5) and *pp* = 1.0 under the k-means partitioning strategy (Supplementary Figure 6). Under maximum-likelihood, the effect is less obvious, as the ML tree under greedy partitioning has *Martialis* sister to formicoids and poneroids but now with only 50% bootstrap support (Supplementary Figure 7). In the k-means maximum-likelihood tree, *Martialis* and Leptanillinae form a clade supported in 99% bootstrap replicates (Supplementary Figure 8).

Removal of GC-rich outgroups (*Ampulex compressa*, *Aporus niger*, *Apterogyna* ZA01, *Chalybion californicum*, and *Chyphotes mellipes*) reinforces support for the topology where Leptanillinae are sister to *Martialis* plus formicoids plus poneroids with *pp* = 1.0 under greedy partitioning (Supplementary Figure 9) and *pp* = 0.98 under k-means in Bayesian analyses (Supplementary Figure 10). The same pattern is present in maximum-likelihood trees, which both show Leptanillinae sister to *Martialis* plus formicoids and poneroids. Under greedy analyses bootstrap support for this clade is 99% (Supplementary Figure 11) and under k-means it is 96% (Supplementary Figure 12).

### 3.4. Compositionally homogeneous matrix

The analyses of homogeneous data partitions result in a tree where *Martialis* is sister to the Leptanillinae with strong support regardless of partitioning scheme and inference method (Table 2). Bayesian trees under both greedy and k-means partitioning strategies show *pp* = 1.00 (Supplementary Figures 13 and 14). Under maximum-likelihood this node is supported in 100% bootstrap replicates under both greedy k-means partitioning strategies (Supplementary Figures 15 and 16).

**Table 2:** Support for selected relationships in Bayesian consensus and maximum-likelihood (ML) trees. Support for Bayesian analyses is expressed in posterior probabilities rounded to two decimal places and for ML in percentage of bootstrap replicates. Ag: Agroecomyrmecinae, Am: Amblyoponinae plus Apomyrminae, for: formicoids, Le: Leptanillinae, Ma: Martialinae, Pa: Paraponerinae, Po: Ponerinae, pon: poneroids, Pr: Proceratiinae. NA signifies a case where the relationship was not recovered in the consensus or maximum-likelihood tree.

| Matrix | Method | Partitioning | Ma + Le | Ma + (for<br>+ pon) | pon<br>monophyl. | Ag + Pa | Po + (Ag<br>+ Pa) | Pr + Am | Pr +<br>(Ag/Pa/Po) | Am +<br>(Ag/Pa/Po) |
| --- | --- | --- | --- | --- | --- | --- | --- | --- | --- | --- |
| Full | ML | greedy | NA | 93 | 100 | 100 | 97 | NA | 77 | NA |
| Full | ML | k-means | 87 | NA | 92 | 98 | 88 | 71 | NA | NA |
| Full | Bayes | greedy | NA | 0.81 | 1.00 | 1.00 | 1.00 | NA | 0.54 | NA |
| Full | Bayes | k-means | 0.99 | NA | 1.00 | 1.00 | 1.00 | 0.59 | NA | NA |
| AT-rich outgr. removed | ML | greedy | NA | 50 | 99 | 100 | 92 | NA | 70 | NA |
| AT-rich outgr. removed | ML | k-means | 99 | NA | 97 | 99 | 83 | 64 | NA | NA |
| AT-rich outgr. removed | Bayes | greedy | 0.91 | NA | 1.00 | 1.00 | 1.00 | 0.5 | NA | NA |
| AT-rich outgr. removed | Bayes | k-means | 1.00 | NA | 1.00 | 1.00 | 1.00 | 0.64 | NA | NA |
| GC-rich outgr. removed | ML | greedy | NA | 99 | 99 | 100 | 95 | NA | 86 | NA |
| GC-rich outgr. removed | ML | k-means | NA | 96 | 75 | 96 | 73 | NA | 45 | NA |
| GC-rich outgr. removed | Bayes | greedy | NA | 1.00 | 1.00 | 1.00 | 1.00 | NA | 0.87 | NA |
| GC-rich outgr. removed | Bayes | k-means | NA | 0.98 | 0.88 | 0.99 | 0.88 | NA | 0.54 | NA |
| Homogeneous | ML | greedy | 100 | NA | 98 | 81 | 99 | NA | NA | 79 |
| Homogeneous | ML | k-means | 100 | NA | 98 | 78 | 100 | NA | NA | 72 |
| Homogeneous | Bayes | greedy | 1.00 | NA | 1.00 | 0.99 | 1.00 | NA | NA | 0.67 |
| Homogeneous | Bayes | k-means | 1.00 | NA | 1.00 | 0.93 | 1.00 | 0.50 | 0.50 | NA |

### 3.5. Simulation

In the maximum-likelihood analyses of simulated alignments imitating first codon positions the *Martialis* plus Leptanillinae clade, i.e., the topology under which the data were simulated, is recovered consistently (Supplementary Figure 17). For the alignments imitating second codon positions *Martialis* emerges as sister to the Leptanillinae in 96 out of 100 trees Supplementary Figure 18), and in the tree derived from the matrix imitating third codon positions *Martialis* is sister to the Leptanillinae in only 64 trees out of 100 (Supplementary Figure 19). The remaining 36 trees show *Martialis* either as sister to the poneroid plus formicoid clade, with Leptanillinae being sister to all other ants, or, alternatively as the sister group to all ants. The combined data set supports the *Martialis* plus Leptanillinae clade in 99 out of 100 trees (Supplementary Figure 20).

### 3.6. Relationships among poneroid subfamillies

The so-called poneroid ant subfamillies that include Agroecomyrmecinae, Amblyoponinae, Apomyrminae, Paraponerinae, Ponerinae, and Proceratiinae form a well supported clade. This result appears more robust to different analytics than the placement of the root of the tree. Support for this clade is often maximum in Bayesian analyses and generally above 90% bootstrap proportion in the maximum-likelihood analyses, except for the data set with GC-rich outgroups removed, where the support is only 75% (Table 2).

Within the poneroid clade, another set of relationships that is well-supported across the analyses is the sister relationship of Agroecomyrmecinae and Paraponerinae, which is significantly supported in all analyses except in the maximum-likelihood trees inferred from the homogeneous data matrix. This clade is in turn sister to the Ponerinae in all analyses, although support varies. This relationship receives maximum support in all Bayesian analyses except for the data matrix with GC-rich outgroups removed. In maximum-likelihood trees support varies between 74% and 97% bootstrap replicates (Table 2).

The most problematic is placement of Proceratiinae and (Amblyoponinae + Apomyrminae), which in some analyses form a clade, and in others form a grade where Amblyoponinae plus Apomyrminae are the sister clade to all other poneroids and Proceratiinae is sister to the remaining subfamilies. The support for both of these alternatives is never significant, however (Table 2).

### 3.7. Non-monophyly of currently recognized genera

Several shallow nodes, well-supported regardless of the data set and analysis method, highlight non-monophyly of genera outside of the formicoid clade (Figure 2; Supplementary Figures 1–16).

In the Leptanillinae, the morphologically derived genus *Anomalomyrma* is nested within *Protanilla*, and two genera known only from males, *Phaulomyrma* and *Yavnella*, are nested within *Leptanilla*.

Within the small subfamily Proceratiinae, the four species of *Proceratium* included in our data matrices were paraphyletic with respect to *Probolomyrmex*.

Under all analyses we recover three non-monophyletic genera within Ponerinae: *Cryptopone gilva* and *Cryptopone hartwigi* included here are only very distantly related, the genus *Euponera* is paraphyletic with respect to the *Cryptopone hartwigi* plus *Fisheropone* clade, and *Mesoponera* is polyphyletic, here represented by *M. melanaria*, which is sister to *Leptogenys*, and *M. ambigua*, here sister to *Streblognathus peetersi*.

### 3.8. Divergence time analyses

Our divergence time analysis recovers a relatively young age for the most common ancestor of crown-group ants, estimated to have lived during the Albian or Aptian ages of the Lower Cretaceous (Figure 3, Table 3; median age 112 Ma, 95% highest posterior density interval 103–123 Ma). The crown formicoids are estimated to have arisen ~101 Ma, closely followed by the split of *Martialis* from the Leptanillinae around 99 Ma, and the origin of poneroids at 92 Ma. The median ages inferred for the subfamilies where our taxon sampling spanned the root node are as follows: 45 for Agroecomyrmecinae, 75 Ma for Ambyloponinae plus Apomyrminae, 55 Ma for Dolichoderinae, 60 Ma for Formicinae, 66 Ma for Leptanillinae, 45 Ma for Myrmeciinae, 61 Ma for Myrmicinae, 73 Ma for Ponerinae, and 65 Ma for Proceratiinae.

**Figure 3:**
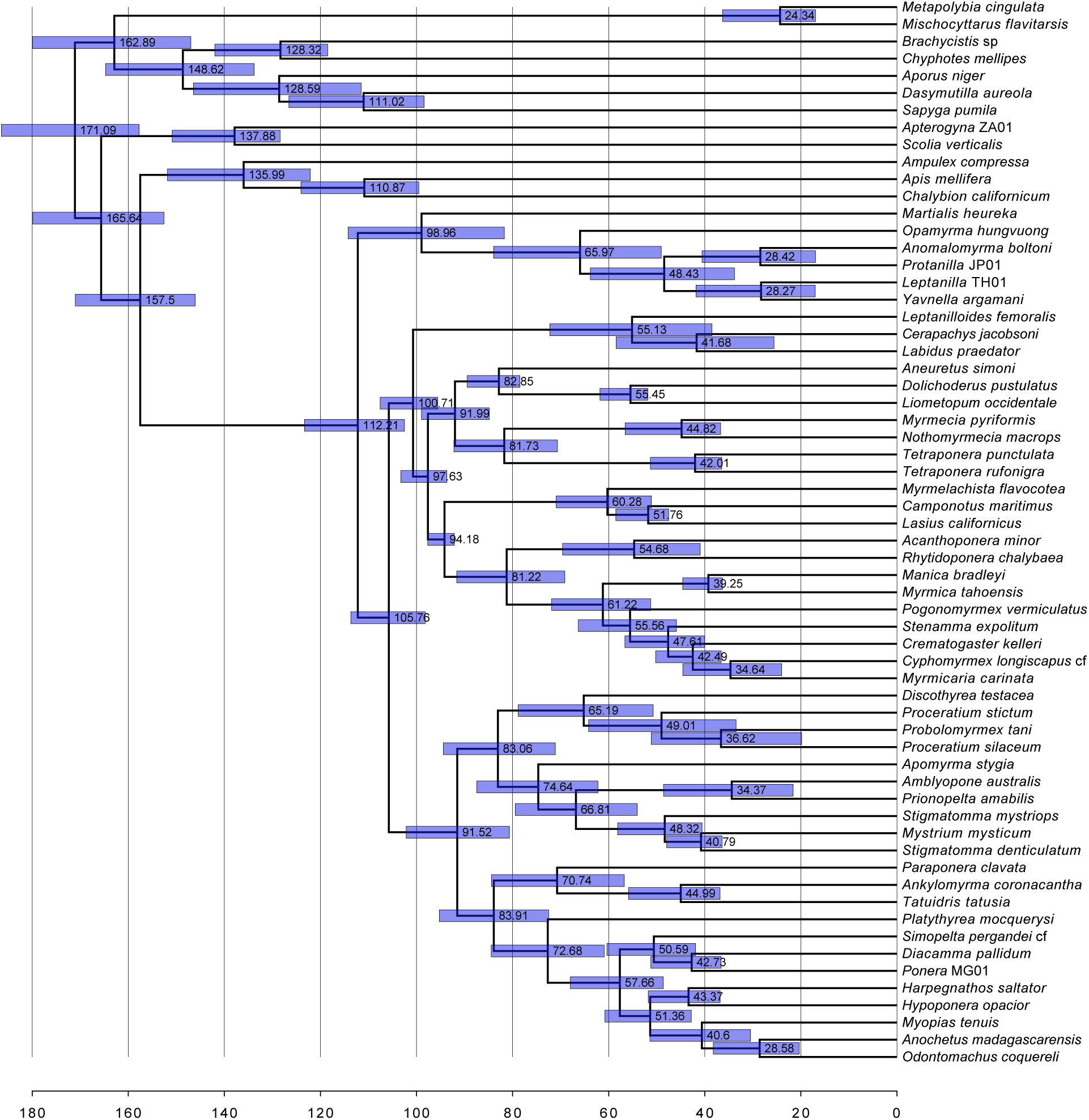
Chronogram from the divergence dating analysis under the fossilized birth-death process with diversified sampling, in MrBayes. Scale is in Ma. Bars depict the 95% highest posterior probability density of each estimate.

**Table 3:**
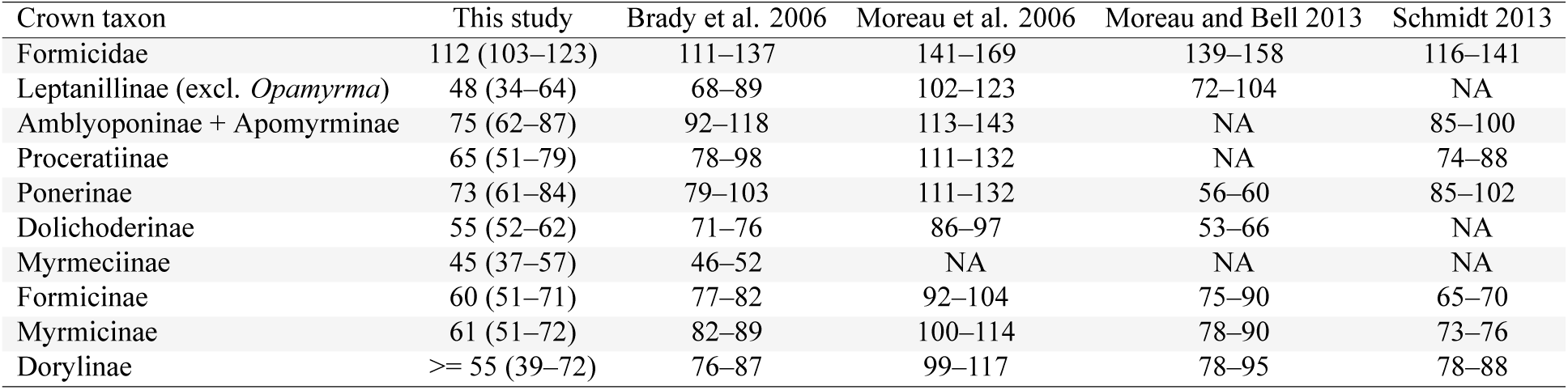
Comparison of divergence time estimates. Numbers for this study referto median node age (95% highest posterior density) in Ma. Numbers from other studies are ranges of means/medians from across all analyses presented.

## 4. Discussion

### 4.1. Compositional heterogeneity and the rooting of the ant tree

Earlier studies recognized the difficulty in rooting the ant tree of life (Brady et al., 2006; Rabeling et al., 2008) and our analyses confirm the supposition (Ward, 2014) that the effects of compositional heterogeneity play a role. The outgroup removal experiments, exclusion of compositionally heterogeneous sites, and simulations all suggest that with greater compositional heterogeneity in the data the abnormally AT-rich Leptanillinae species are drawn more strongly to the base of the ant tree. As a result of this, the more GC-rich *Martialis* can erroneously cluster with the clade of formicoids and poneroids, as in Kück et al. (2011). When compositional heterogeneity is accounted for, as in the homogeneous data matrix, the Leptanillinae and *Martialis* emerge as sister taxa forming a strongly supported clade that is sister to all other ants.

The selective outgroup removal experiment shows a trend in which support for the AT-rich Leptanillinae as sister to all other ants including *Martialis* is the strongest when only AT-rich outgroups are retained, moderate or weak when all outgroups (those that are AT- and GC-rich) are retained, and the weakest when all the AT-rich outgroups are removed. This suggests that the AT content of outgroup taxa indeed causes attraction of the above-average AT-rich Leptanillinae to the base of the ant tree. This finding has interesting implications for outgroup choice in phylogenetics in general, as it suggests that choice of outgroups should be made not only on the basis of their close relationship to the ingroup taxa, but also taking into account sequence divergence and other properties relative to the ingroup (Takezaki and Nishihara, 2016)

Finally, our simulations show that compositional heterogeneity similar to that in our empirical data matrix has the potential to cause bias resulting in the incorrect placement of Leptanillinae as sister to *Martialis* plus poneroids plus formicoids. The matrix designed to imitate third codon positions was simulated on a tree where *Martialis* was sister to Leptanillinae, but the maximum-likelihood trees inferred from this alignment often recover *Martialis* sister to all other ants or sister to the poneroids plus formicoids, both topologies recovered in previous studies (Rabeling et al., 2008; Kück et al., 2011). The tree topology used for the simulation is correctly recovered from the alignments imitating first and second codon positions in most cases. These alignments were simulated with base frequencies drawn from our empirical alignment and under a tree-homogeneous model. On a combined simulated data set the negative effect of sites emulating third codon positions is overwhelmed by the homogeneous data and the inferred tree is consistent with the topology on which the data were simulated in 99 out of 100 maximum-likelihood trees obtained from the simulated concatenated data. These effects appear not to be as strong as seen in our empirical data, but the simulation is a simplistic scenario that is likely to involve fewer confounding factors. In particular, 1) the same substitution model that generated the data could be used for inference (minus compositional heterogeneity), 2) the simulations attempted to capture only one dimension of the process heterogeneity, and 3) in our empirical data matrix compositionally heterogeneous sites were actually over-represented compared to the simulated matrix because of the heterogeneous *28S* partition, which was not taken into account when simulating the matrix imitating the non-homogenous third codon positions. The results from simulations imitating compositionally heterogeneous third codon position data demonstrate that compositional heterogeneity, at least in principle, has the potential to impact the position of *Martialis*.

If our interpretation of these results is correct, the species-poor clade of blind and subterranean Martialinae and Leptanillinae is the sister group to the remaining 99.5% species of the Formicidae. As pointed out before (Brady et al., 2006; Rabeling et al., 2008; Pie and Feitosa, 2015), this does not necessarily mean that the most-recent common ancestor of the ants was blind and lived underground (Lucky et al., 2013). Rather, this fact may reflect lower relative extinction rates experienced by these ants, perhaps due to the long-term stability of their subterranean environments, or different relative probabilities of evolutionary transitions between subterranean and epigaeic habits.

### 4.2. Relationships of poneroid subfamillies

In addition to further insight into the placement of the root of the ant phylogeny, we find evidence for poneroid monophyly. The question of poneroid monophyly vs. paraphyly was the second outstanding issue in higher ant phylogeny highlighted in a recent review (Ward, 2014). All our empirical analyses suggest poneroid monophyly, and in most instances this clade receives significant support, with the notable exception of analyses of the data matrix from which GC-rich outgroups were removed, i.e., where the phylogeny was potentially more susceptible to bias. Poneroid monophyly was first recovered in Moreau et al. (2006) but this result was questioned as doubtful by Brady et al. (2006), who emphasized contradictory results from their Bayesian and maximum-likelihood analyses of which the former supported monophyletic poneroids but the latter did not. Brady et al. (2006) also conducted ingroup-only analyses which supported topologies where no possible rooting could result in poneroid monophyly. Although we did not perform ingroup-only analyses here, taking into account more comprehensive taxon sampling, the higher amount of sequence data, and insensitivity of poneroid monophyly to the different data treatments, we interpret the support for the poneroid clade as strong. Poneroid monophyly has also been recovered in other phylogenetic studies of ants, including Moreau et al. (2006), some analyses of Brady et al. (2006), Ward and Fisher (2016), and a phylogenomic study (Branstetter et al., 2017b).

Similarly strongly-supported results involve the relationships among the poneroid clade sub-families Agroecomyrmecinae, Paraponerinae, and Ponerinae. Although very disparate morphologically, Agroecomyrmecinae emerge as sister to Paraponerinae with significant support in all analyses, and this clade is in turn sister to the Ponerinae.

Our data are inconclusive on the relative position of Proceratiinae and (Amblyoponinae plus Apomyrminae), which sometimes emerge as a clade and at other times as a grade relative to the clade composed of Agroecomyrmecinae, Paraponerinae, and Ponerinae. If other results presented here are confirmed, the placement of (Ambyloponinae plus Apomyrminae) and Proceratiinae within the poneroid group would remain the last unsolved subfamily-level relationship in ants.

### 4.3. Non-monophyly of currently recognized genera

Our analyses recover several of the currently recognized ant genera as para- or polyphyletic. Although we do not favor inclusion of non-monophyletic groups in a classification, here we are only highlighting existing problems without proposing any formal taxonomic changes. We feel that proposing satisfactory resolutions requires additional research, as explained for each case below.

Among the Leptanillinae, we find the morphologically derived genus *Anomalomyrma* nested within samples identified by us as *Protanilla*. A more comprehensive evaluation of *Protanilla*-like leptanillines, including both males and workers, should be carried out for a better understanding of diversity within the group. We find two other leptanilline genera, *Phaulomyrma* and *Yavnella*, nested within *Leptanilla*. Both these genera were described based on males not associated with workers (Wheeler and Wheeler, 1930; Kugler, 1987). Because the characters defining and differentiating leptanilline lineages based on males are not well understood (Ogata et al., 1995) and all of our *Leptanilla* specimens were males, we feel it would be premature to propose taxonomic changes. A critical reappraisal of leptanilline taxonomy using both morphology and molecular phylogenetics is clearly needed.

Our analyses find the proceratiine genus *Probolomyrmex* nested within the larger genus *Proceratium*. Several species currently in *Proceratium* were classified in the erstwhile genus *Sysphingta*. The differentiation between the two taxa was mostly based on the structure of the clypeus and shape of the petiole. Based on these characters, two of the *Proceratium* species included in our phylogeny, *P. avium* and *P. stictum*, would fit the old concept of *Sysphingta*, while the two other, *Proceratium silaceum* and *Proceratium* SC02, match *Proceratium* sensu stricto (*P. silaceum* is the type species of the genus). Previous authors (Brown, 1958; Urbani and Andrade, 2003), however, showed that considerable variation exists with regard to the characters originally used to distinguish *Proceratium* from *Sysphingta. Proceratium* taxonomy would thus benefit from a focused study and a re-evaluation of morphology under a modern phylogenetic framework.

Despite recent comprehensive taxonomic and phylogenetic work focusing on the Ponerinae (Schmidt, 2013; Schmidt and Shattuck, 2014), our analyses reveal three non-monophyletic genera within the subfamily.

*Cryptopone* is a case of a polyphyletic genus. The two species included here, *C. gilva* and *C. hartwigi*, are only very distantly related. The former is a part of the *Ponera* genus-group and the latter sister to *Fisheropone* and a part of the *Odontomachus* genus-group, as defined by Schmidt (2013). As noted by Schmidt and Shattuck (2014), the resolution of *Cryptopone* taxonomy would require a more thorough revision and sampling of all species attributed to this genus. Notably, our phylogeny did not include any species placed in the erstwhile genus *Wadeura*, which may well turn out to be yet another lineage unrelated to the type species *C. testacea*.

*Euponera*, which was recognized to form two morphologically distinct species groups by Schmidt and Shattuck (2014), is represented by *E. brunoi* and *E. sikorae* in our data set. In our phylogeny, *E. brunoi* is more closely related to ”*Cryptopone*” *hartwigi* and *Fisheropone* than *E. sikorae*. Together, the paraphyletic *Euponera*, the species ”*Cryptopone*” *hartwigi*, and *Fisheropone ambigua* form a well-supported group with well-resolved internal relationships. *Euponera* is divisible into two groups based on morphology but there are several species that cannot be placed with certainty even within the genus as presently defined (Schmidt and Shattuck, 2014). Assignment of *Euponera* species into two different genera should thus be postponed until more evidence is available.

*Mesoponera*, also found to be polyphyletic in our analyses, presents a particularly taxonomically challenging genus that would require a more comprehensive reexamination of morphology and inclusion of more species in a phylogeny for satisfactory resolution. See Schmidt and Shattuck (2014) for a more thorough discussion.

### 4.4. The age of extant ants

Our divergence-time analysis indicates that the most recent common ancestor of living ants originated during the Lower Cretaceous (103–124 Ma; median 113 Ma), an age estimate considerably younger than those obtained by some other recent studies. Moreau et al. (2006) concluded that ants most likely arose 140–169 Ma while Moreau and Bell (2013) arrived at an estimate of 139–158 Ma. In contrast, Brady et al. (2006) proposed a younger age for the crown ants, 116–133 Ma.

Our study parallels the pattern for age estimates of the order Hymenoptera, which has often been inferred to be much older than the oldest hymenopteran fossils (Ronquist et al., 2012; Peters et al., 2017), but under fossilized birth-death process with diversified sampling its estimated age fell to within 20 Ma from the oldest known fossils (Zhang et al., 2016). In our study, the median age for the crown Formicidae, at 113 Ma, is also about 20 Ma older than the oldest undisputed crown-group fossils (Grimaldi and Agosti, 2000; Barden, 2017).

The ages we recovered for ant subfamilies where either few old crown-group fossils are known or only a few taxa were sampled are almost certainly underestimated (e.g. Dolichoderinae at 55 Ma, cf. Ward et al. (2010); Formicinae at 60 Ma, cf. Blaimer et al. (2015)). Future studies including more genus-level sampling of extant and extinct taxa are likely to modify these estimates in the direction of older dates.

## 5. Concluding remarks

Although more sequence data have often been shown to help resolve difficult phylogenetic questions, our study of ant phylogeny shows that systematic bias not accounted for by the commonly used tree-homogeneous models may adversely affect phylogenetic inference. Simply increasing the amount of data can in fact be detrimental if added sequences have properties that violate model assumptions (Huelsenbeck and Hillis, 1993), such as the substantial among-taxon compositional heterogeneity present in third codon positions and ribosomal 28S in our data set. The ideal solution to this problem would be use of substitution models that take into account process heterogeneity across the tree. Such models have been proposed (Foster, 2004; Blanquart and Lartillot, 2008; Jayaswal et al., 2014), but unfortunately their current implementations do not scale well for larger data sets, even for the modest amount of data present in our alignment. Alternatively, one can assess model adequacy through simulation-based tests of compositional heterogeneity (Foster, 2004), as in the current study.

The phylogenetic hypothesis presented here for deep nodes of the ant tree of life will soon be tested with genomic-scale data. Recent advances in sequencing and analysis has already produced data matrices with hundreds or even thousands of loci (Faircloth et al., 2014; Blaimer et al., 2015; Branstetter et al., 2017b). As the amount of available sequence data increases, it is important that the potential for model violation is carefully evaluated, as large data sets will likely be less prone to uncertainty but instead may give strongly-supported results that are wrong instead (Philippe and Roure, 2011). Tests of compositional heterogeneity, posterior predictive approaches to assessing model fit (Bollback, 2002; Brown and ElDabaje, 2008; Doyle et al., 2015), or sensitivity of results to removal of sites likely to introduce bias (Goremykin et al., 2015) should become a part of the standard phylogenomics toolkit.

Our understanding of the timeline of ant evolution will also likely benefit from more biologically realistic models resulting from recent developments in divergence-dating, such as placement of fossils using explicit information about morphology (Ronquist et al., 2012; Zhang et al., 2016), and from inclusion of more sequence data as well as more comprehensive taxon sampling.

## 6. Acknowledgements

For providing specimens used in this study we are grateful to Bob Anderson, Himender Bharti, Lech Borowiec, Bui Tuan Viet, Jim Carpenter, Katsuyuki Eguchi, Juergen Gadau, Dave General, Benoit Guénard, Nihara Gunawardene, Peter Hawkes, Armin Ionesco-Hirsch, Bob Johnson, Yeo Kolo, John LaPolla, John Lattke, Jack Longino, Randy Morgan, Michael Ohl, Christian Peeters, James Pitts, Julian Resasco, Keve Ribardo, Riitta Savolainen, Chris Schmidt, Justin Schmidt, Mike Sharkey, Andy Suarez, Simon van Noort, and Masashi Yoshimura. This research was supported by U.S. National Science Foundation grants EF-0431330, DEB-1402432, DEB-0842204, DEB-1354996, DEB-1456964, and DEB-1654829.

---

**Supplementary Figure 1:**
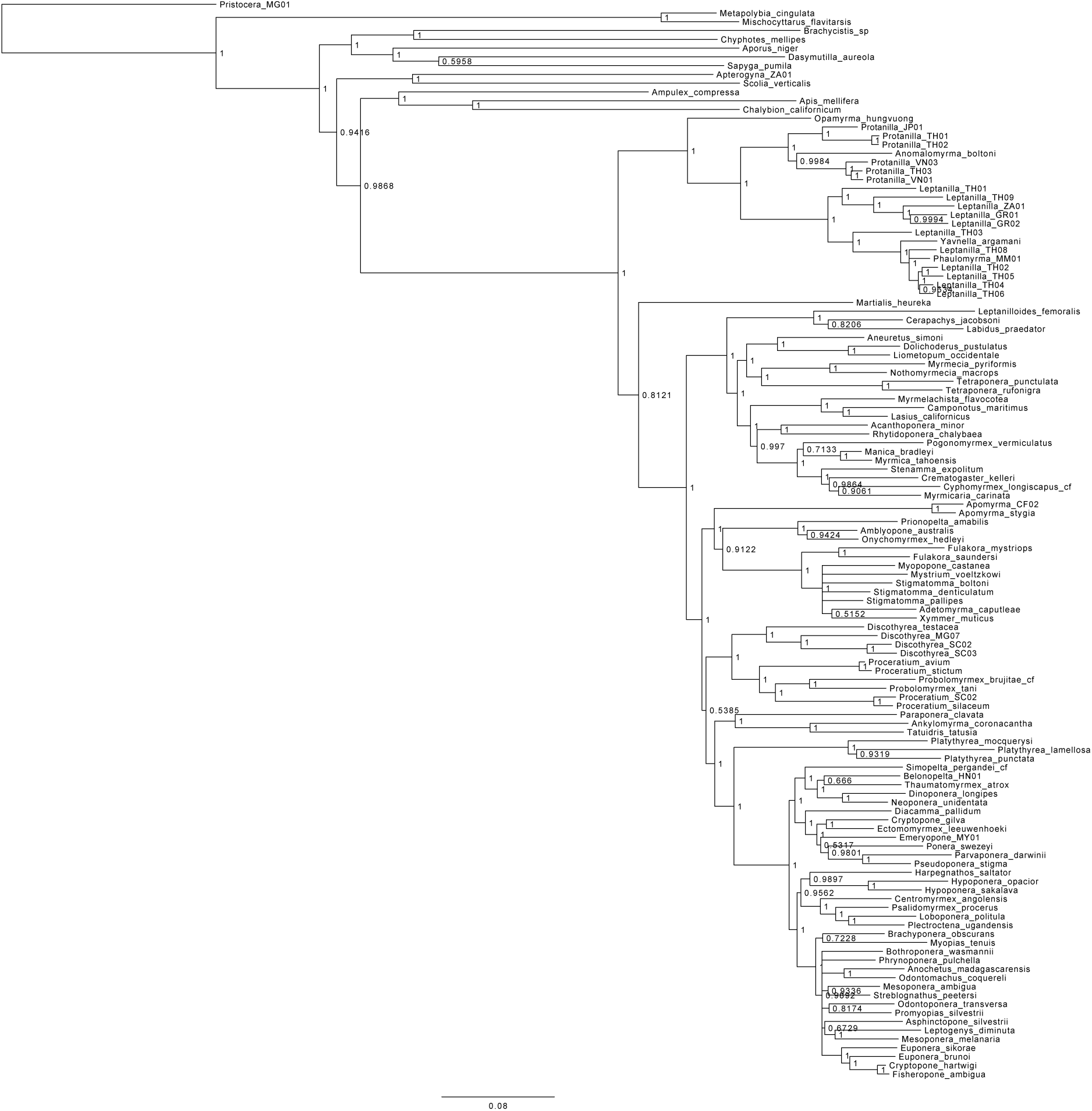
Bayesian consensus tree inferred under greedy partitioning strategy for full data set.

**Supplementary Figure 2:**
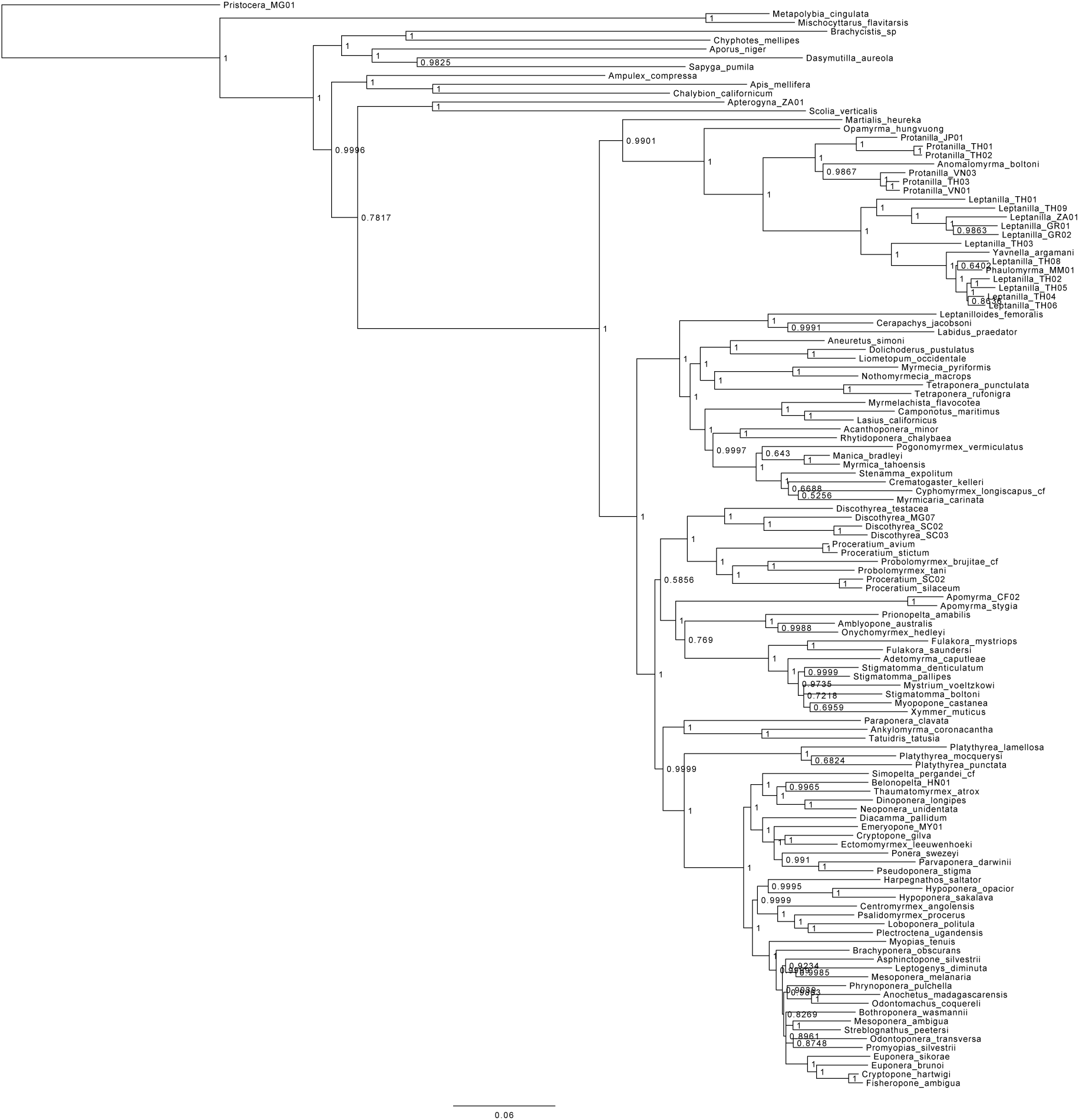
Bayesian consensus tree inferred under k-means partitioning strategy for full data set.

**Supplementary Figure 3:**
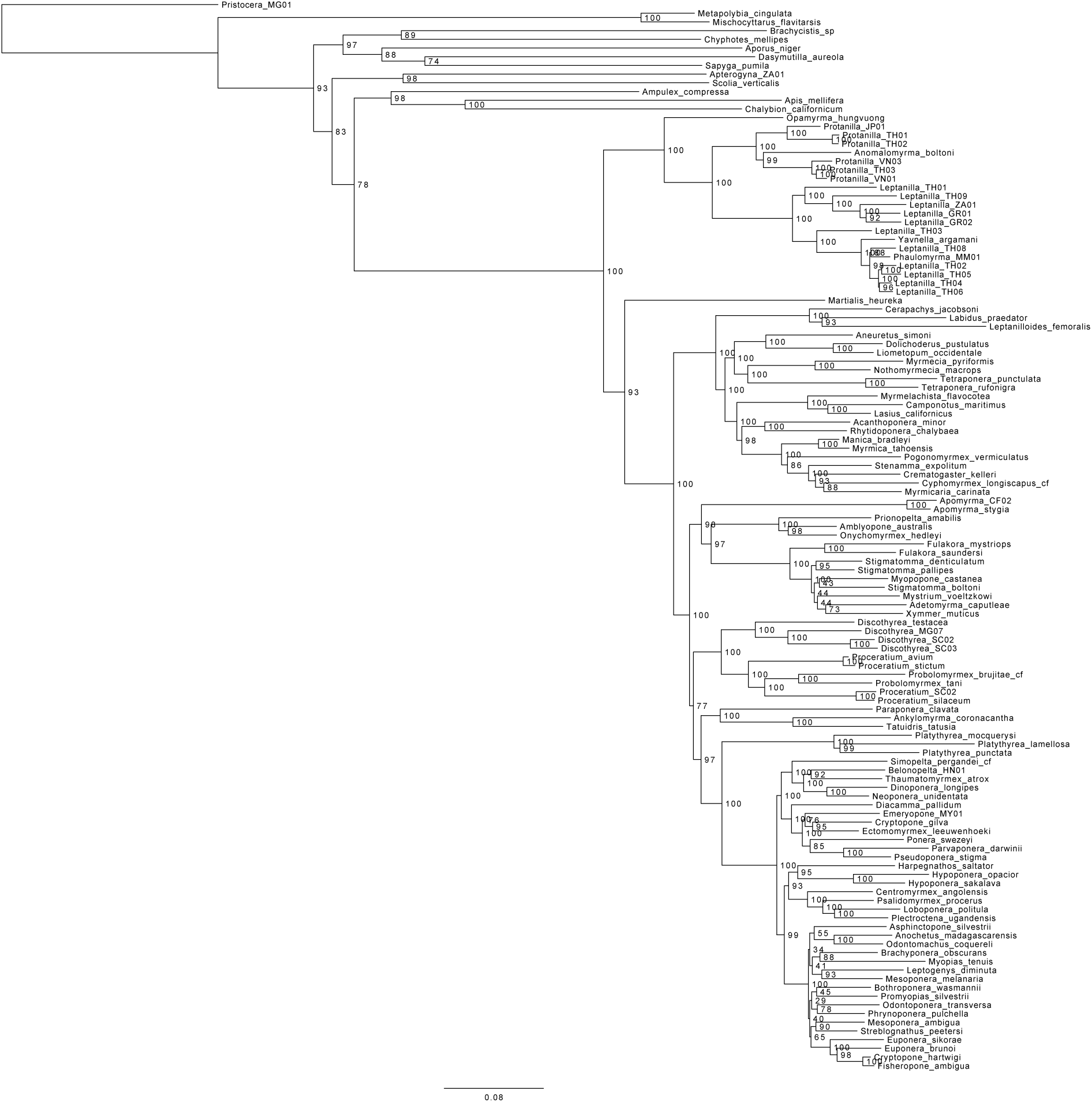
Maximum-likelihood tree inferred under greedy partitioning strategy for full data set.

**Supplementary Figure 4:**
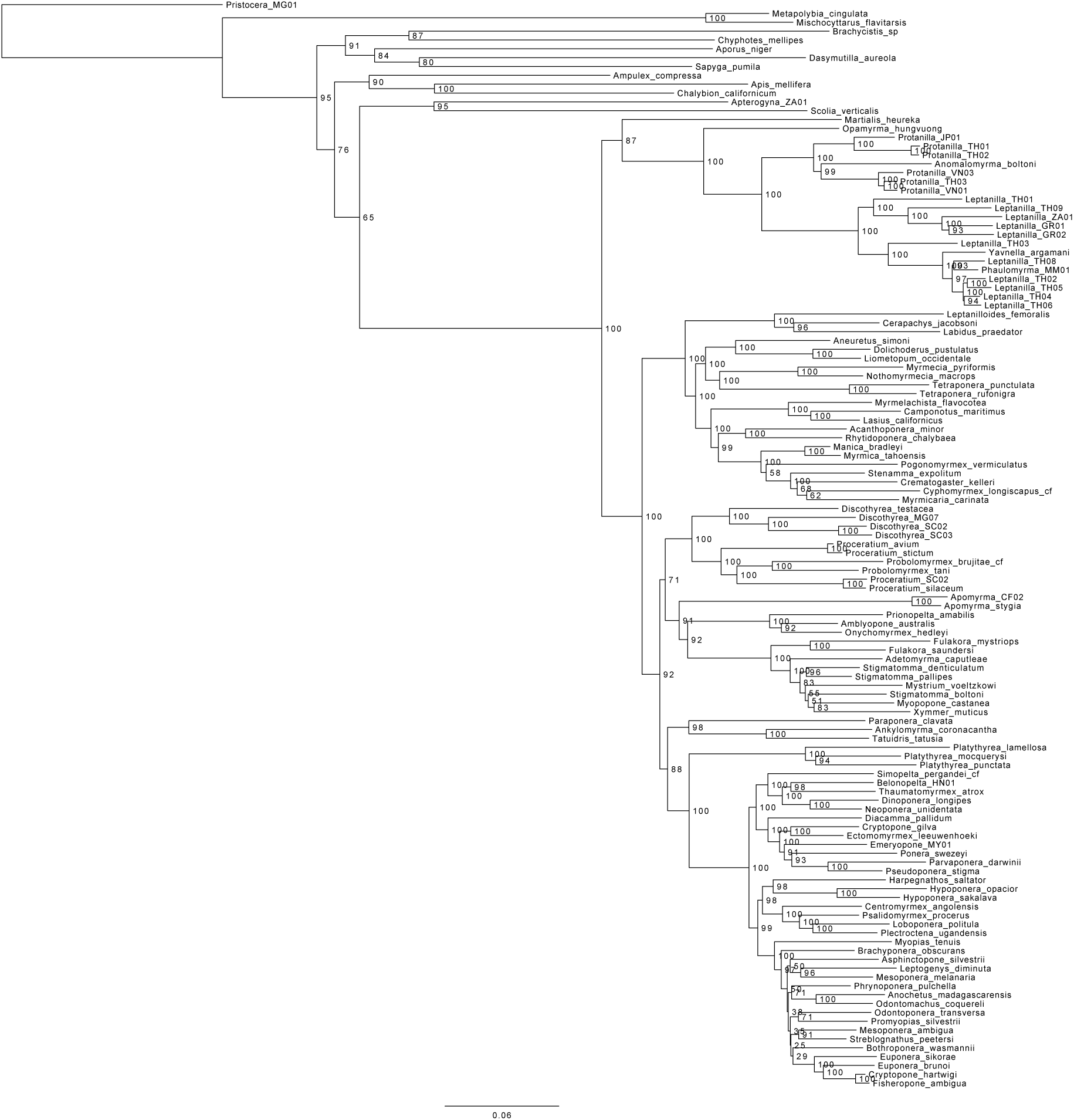
Maximum-likelihood tree inferred under k-means partitioning strategy for full data set.

**Supplementary Figure 5:**
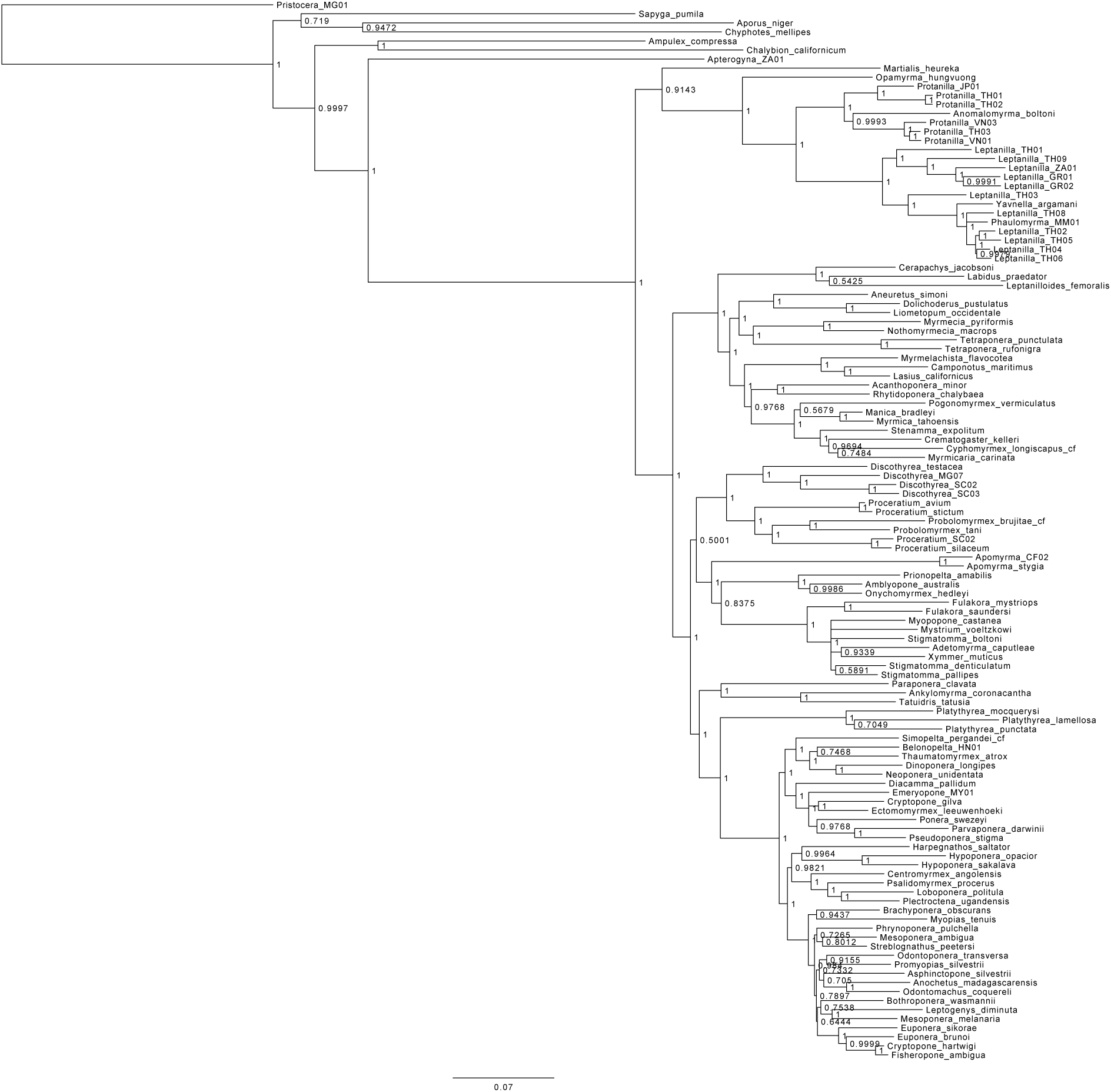
Bayesian consensus tree inferred under greedy partitioning strategy for datset with AT-rich outgroups removed.

**Supplementary Figure 6:**
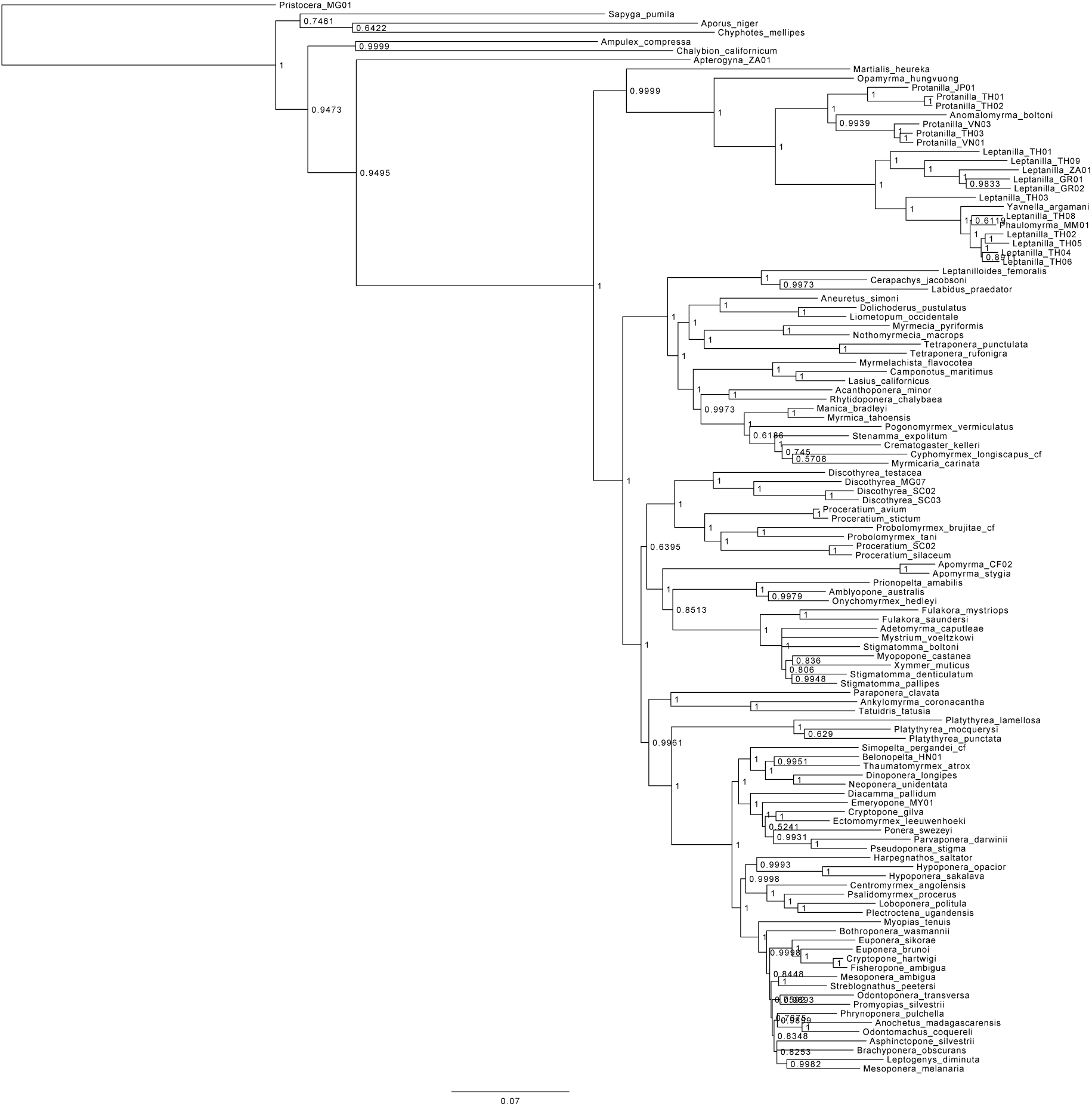
Bayesian consensus tree inferred under k-means partitioning strategy for datset with AT-rich outgroups removed.

**Supplementary Figure 7:**
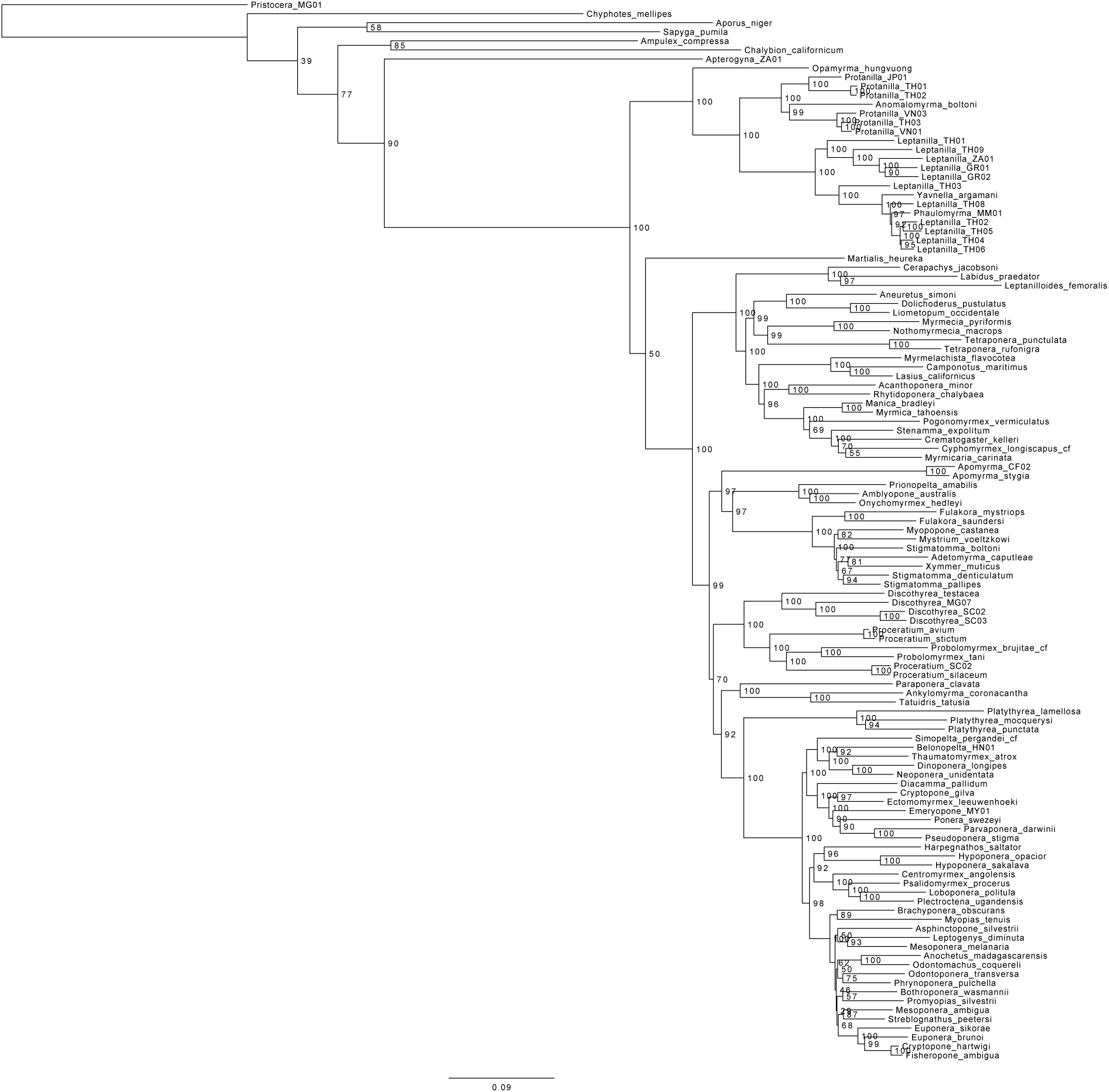
Maximum-likelihood tree inferred under greedy partitioning strategy for datset with AT-rich outgroups removed.

**Supplementary Figure 8:**
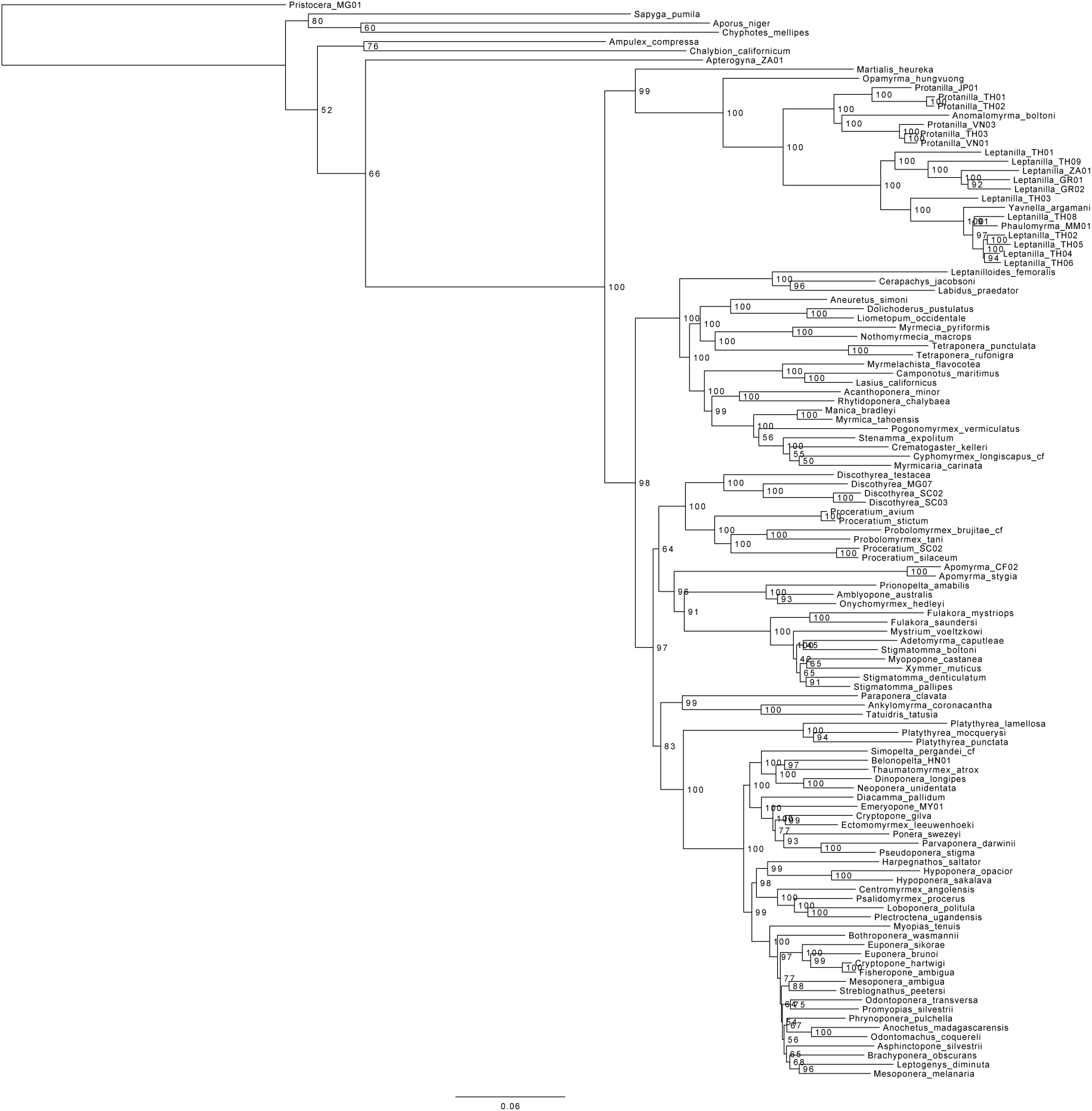
Maximum-likelihood tree inferred under k-means partitioning strategy for datset with AT-rich outgroups removed.

**Supplementary Figure 9:**
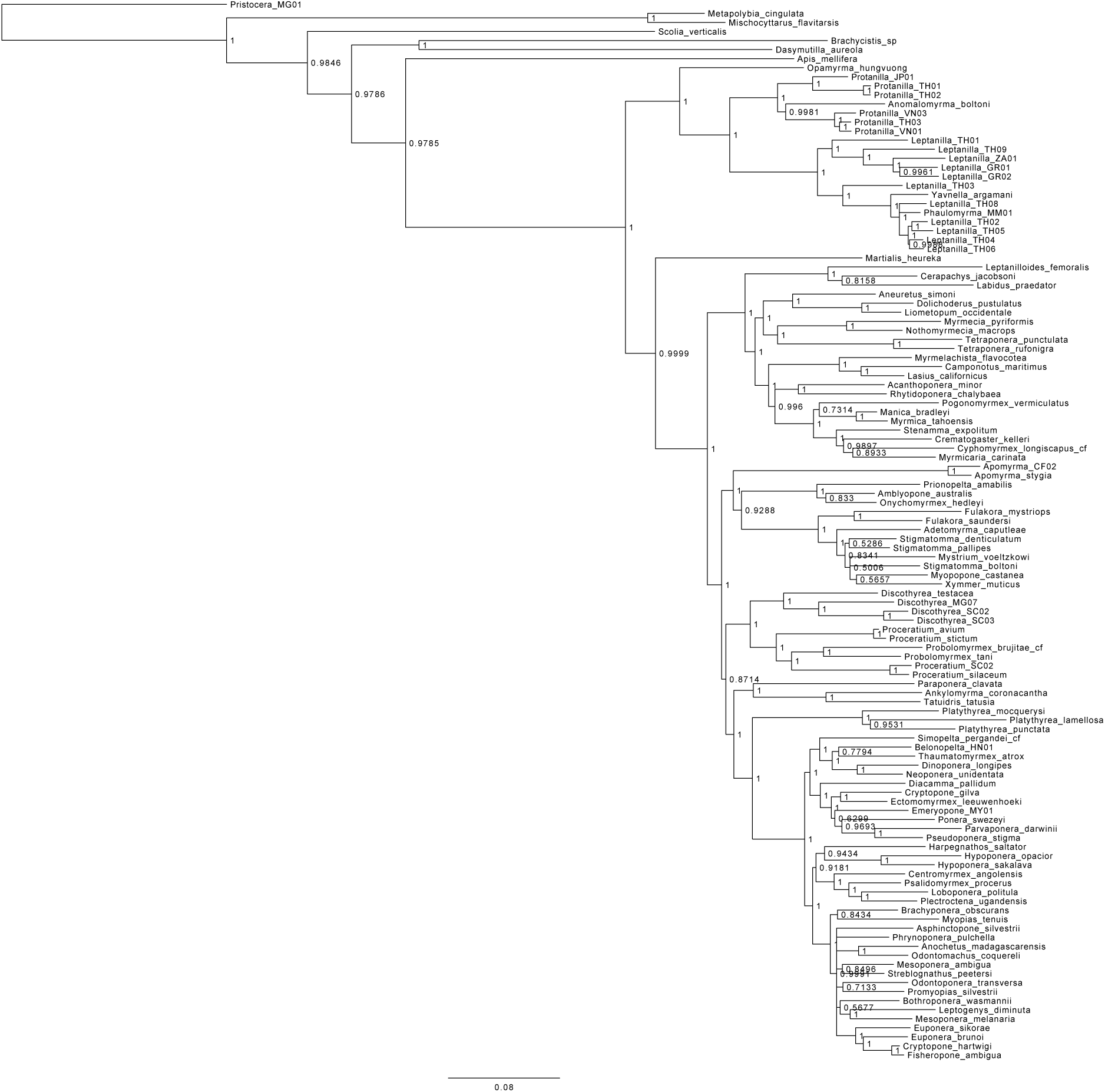
Bayesian consensus tree inferred under greedy partitioning strategy for datset with GC-rich outgroups removed.

**Supplementary Figure 10:**
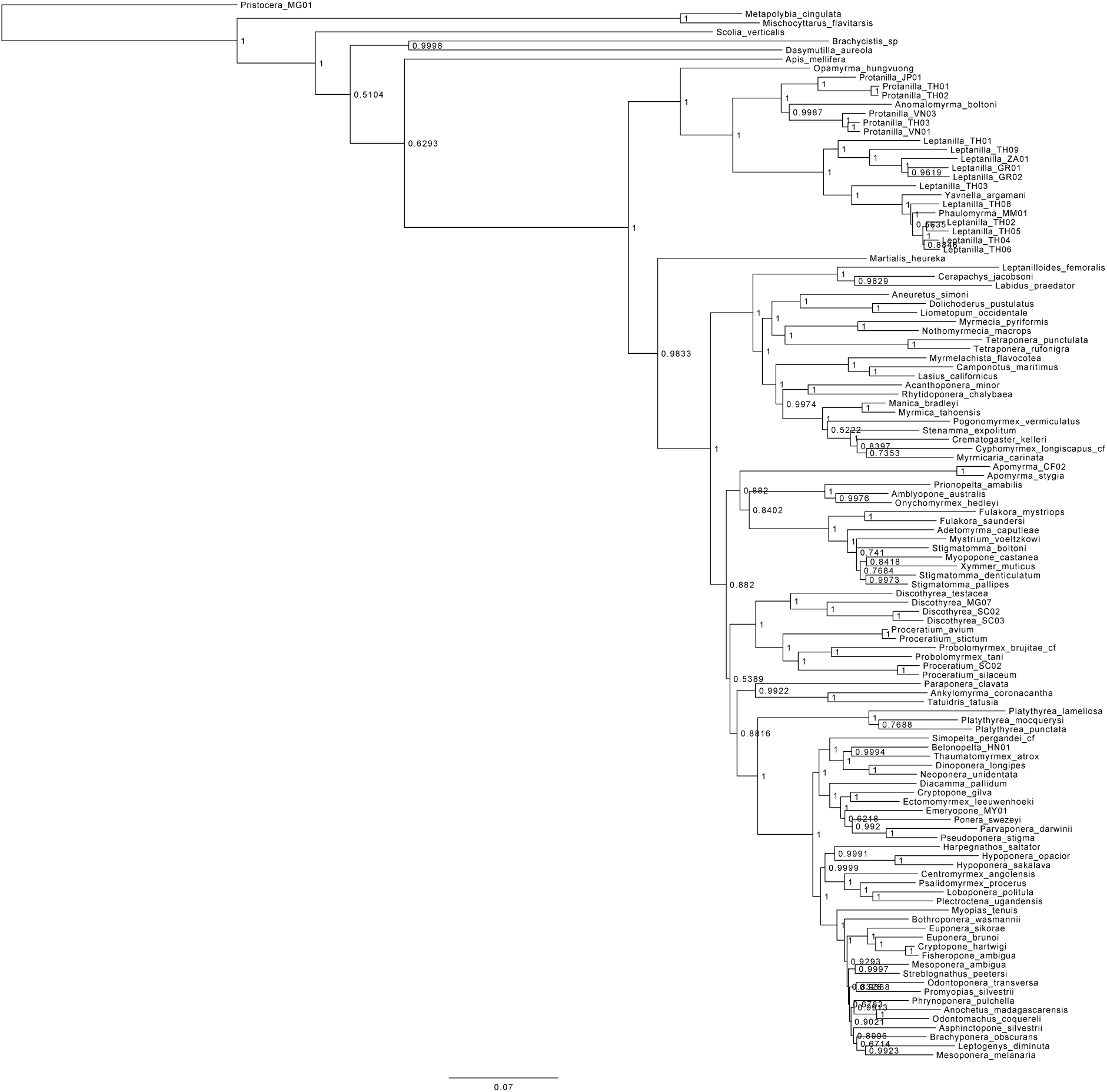
Bayesian consensus tree inferred under k-means partitioning strategy for datset with GC-rich outgroups removed.

**Supplementary Figure 11:**
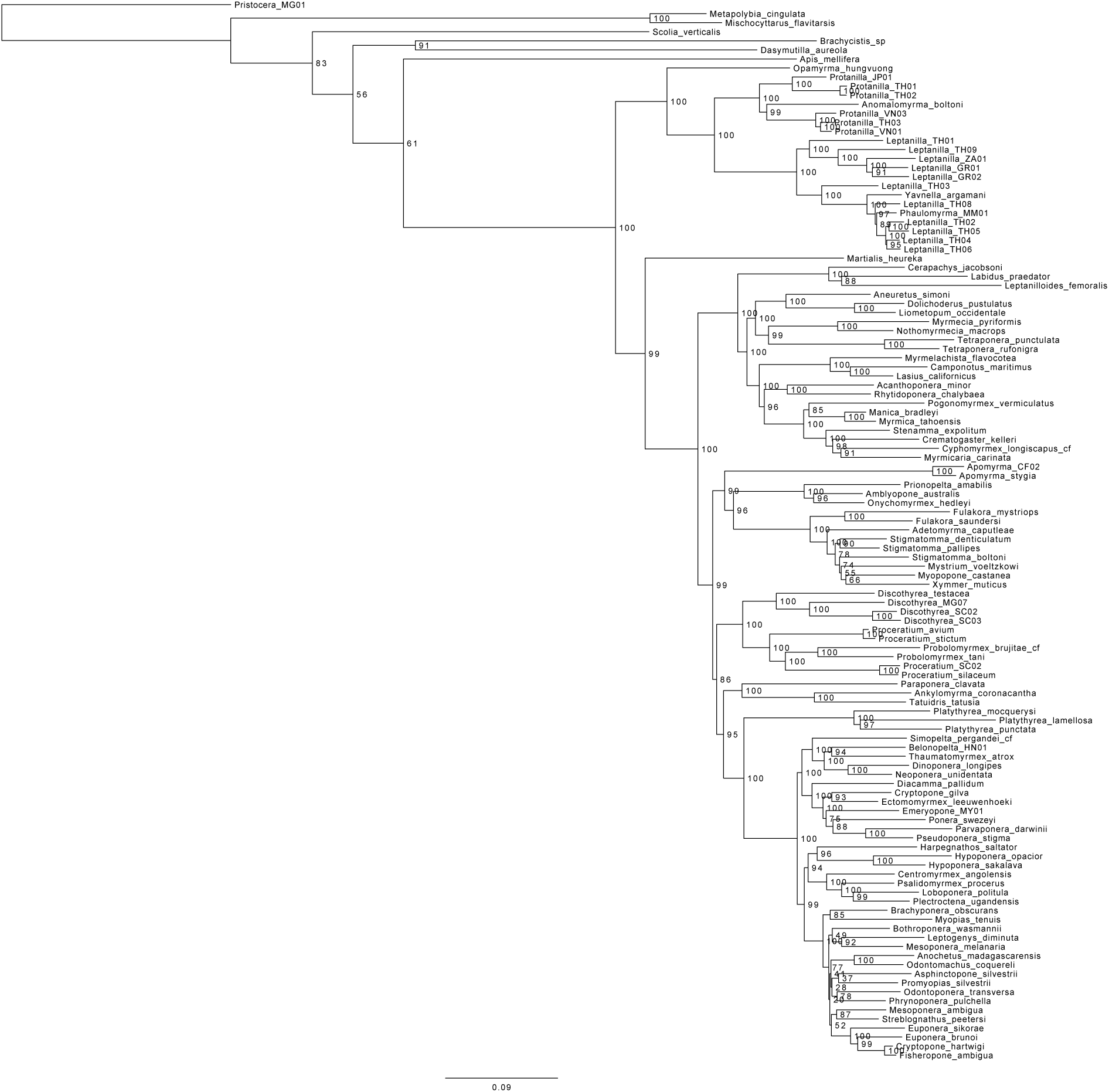
Maximum-likelihood tree inferred under greedy partitioning strategy for datset with GC-rich outgroups removed.

**Supplementary Figure 12:**
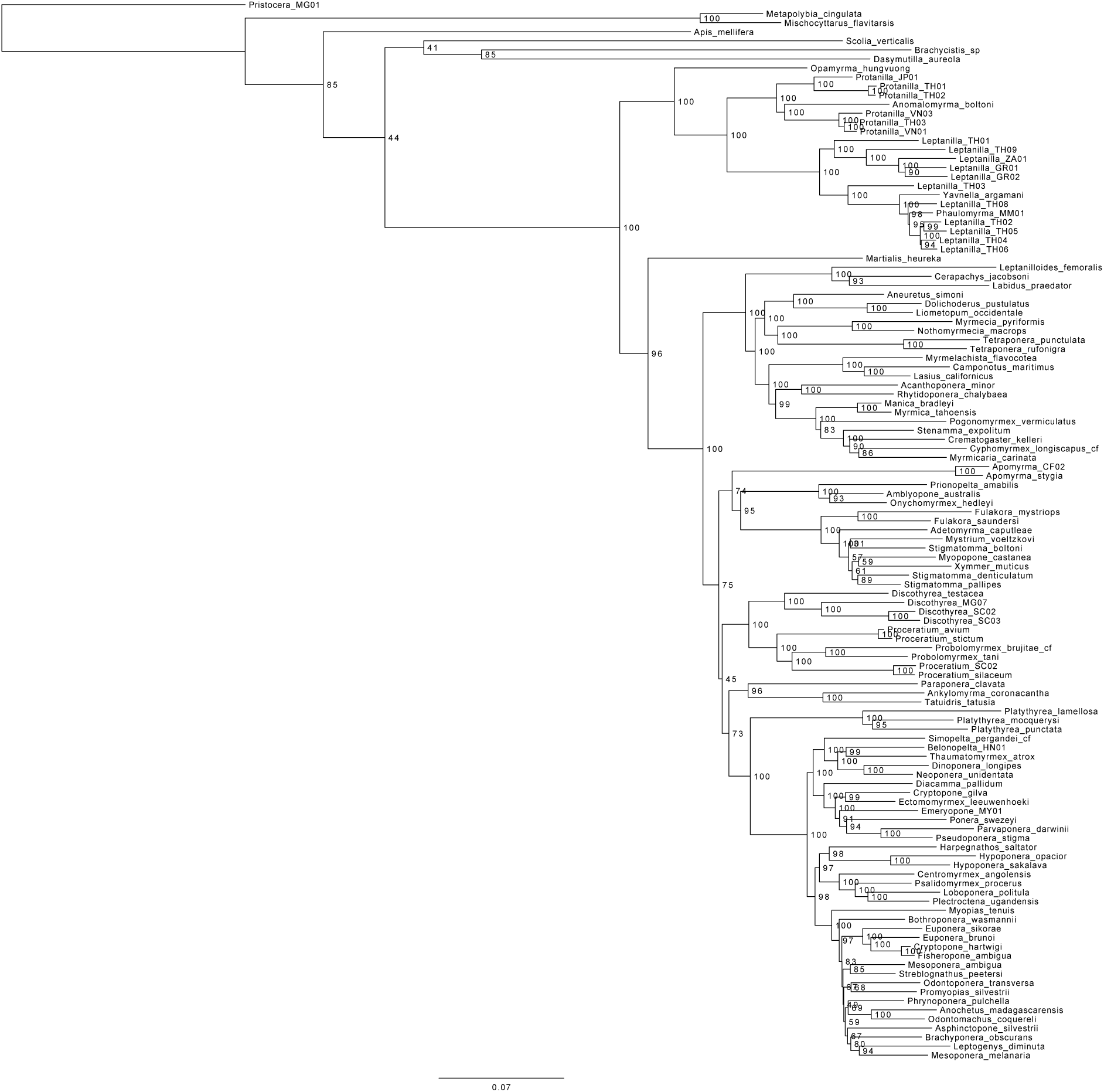
Maximum-likelihood tree inferred under k-means partitioning strategy for datset with GC-rich outgroups removed.

**Supplementary Figure 13:**
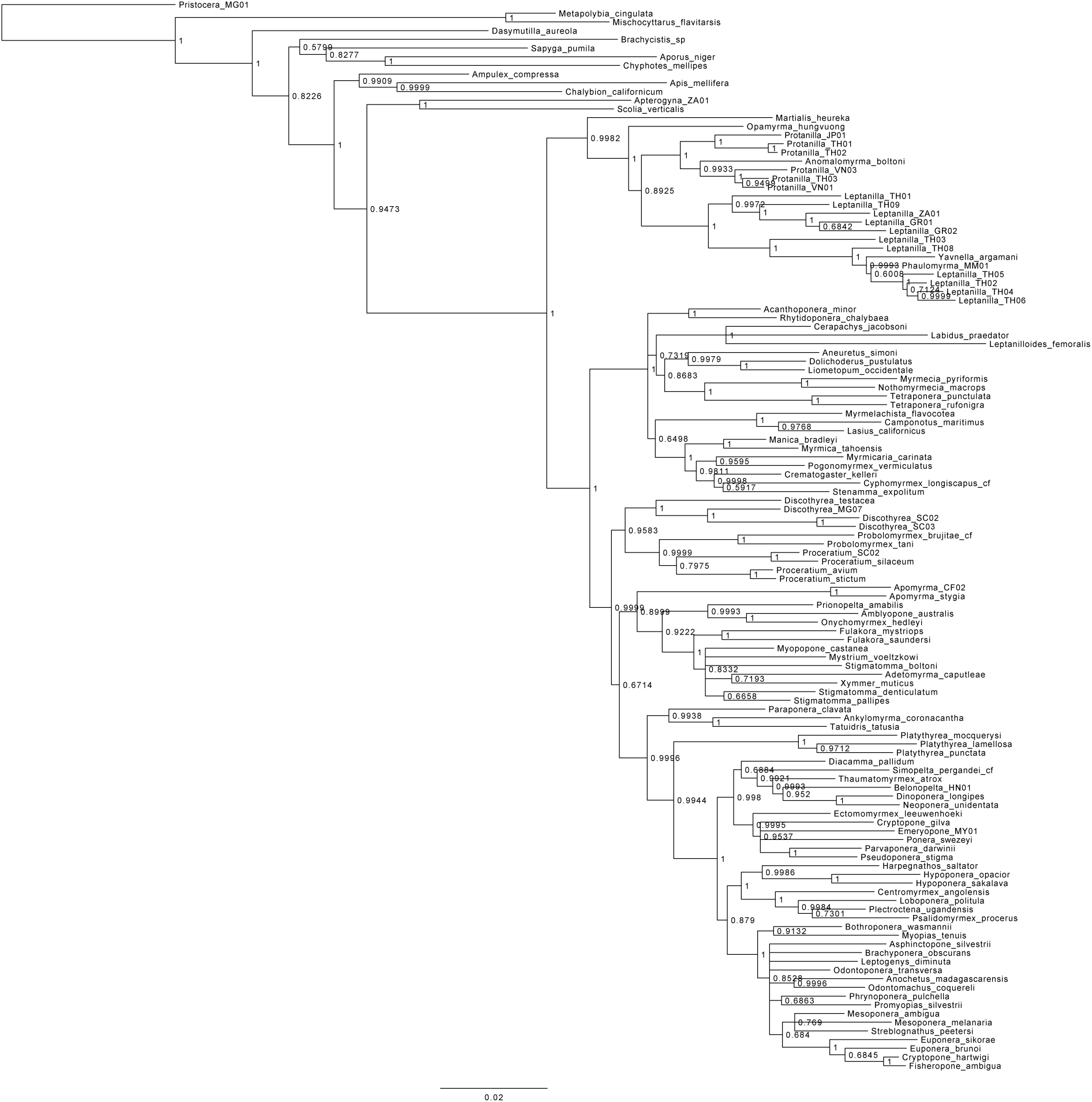
Bayesian consensus tree inferred under greedy partitioning strategy for compositionally homogeneous datset.

**Supplementary Figure 14:**
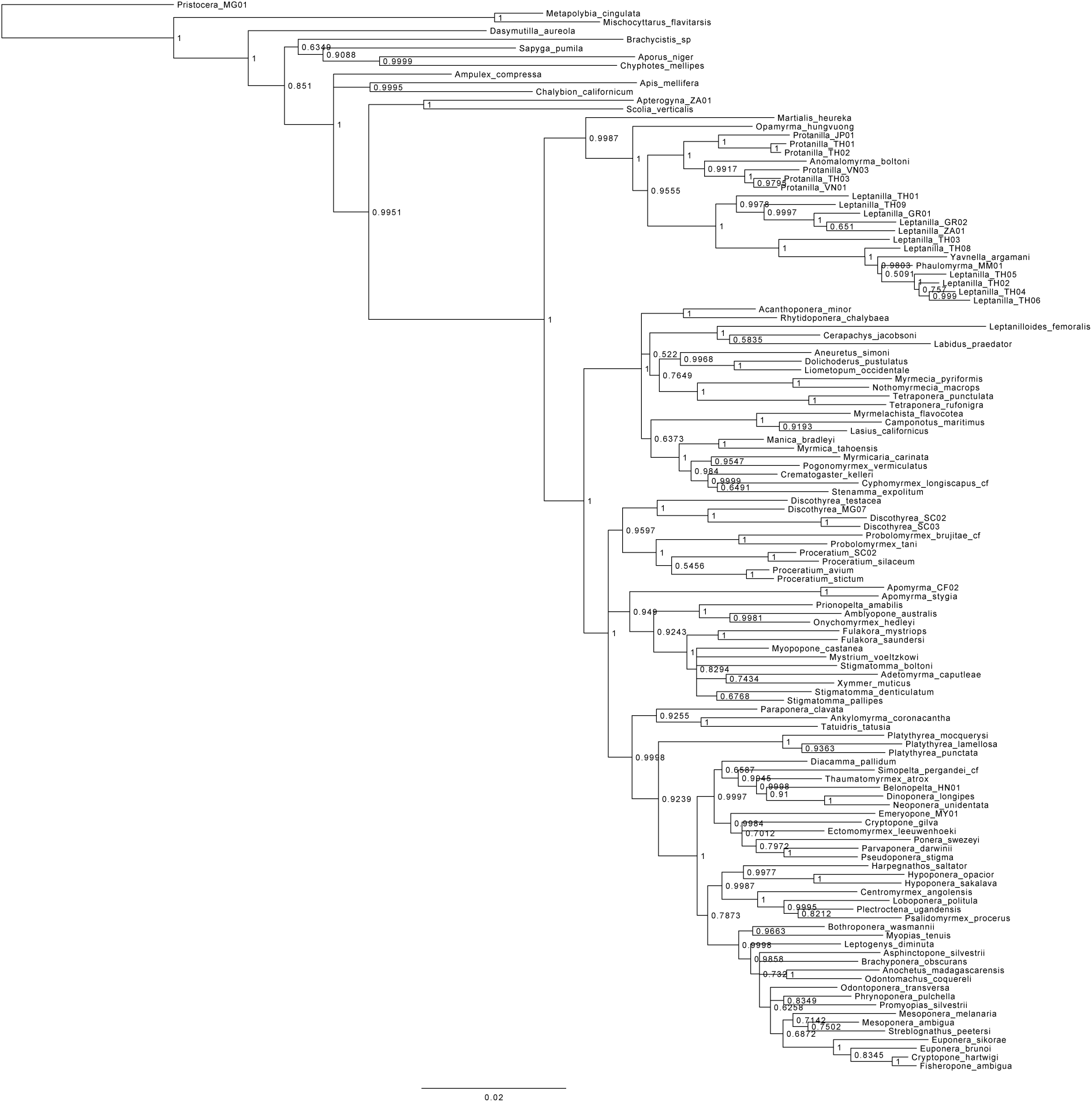
Bayesian consensus tree inferred under k-means partitioning strategy for compositionally homogeneous datset.

**Supplementary Figure 15:**
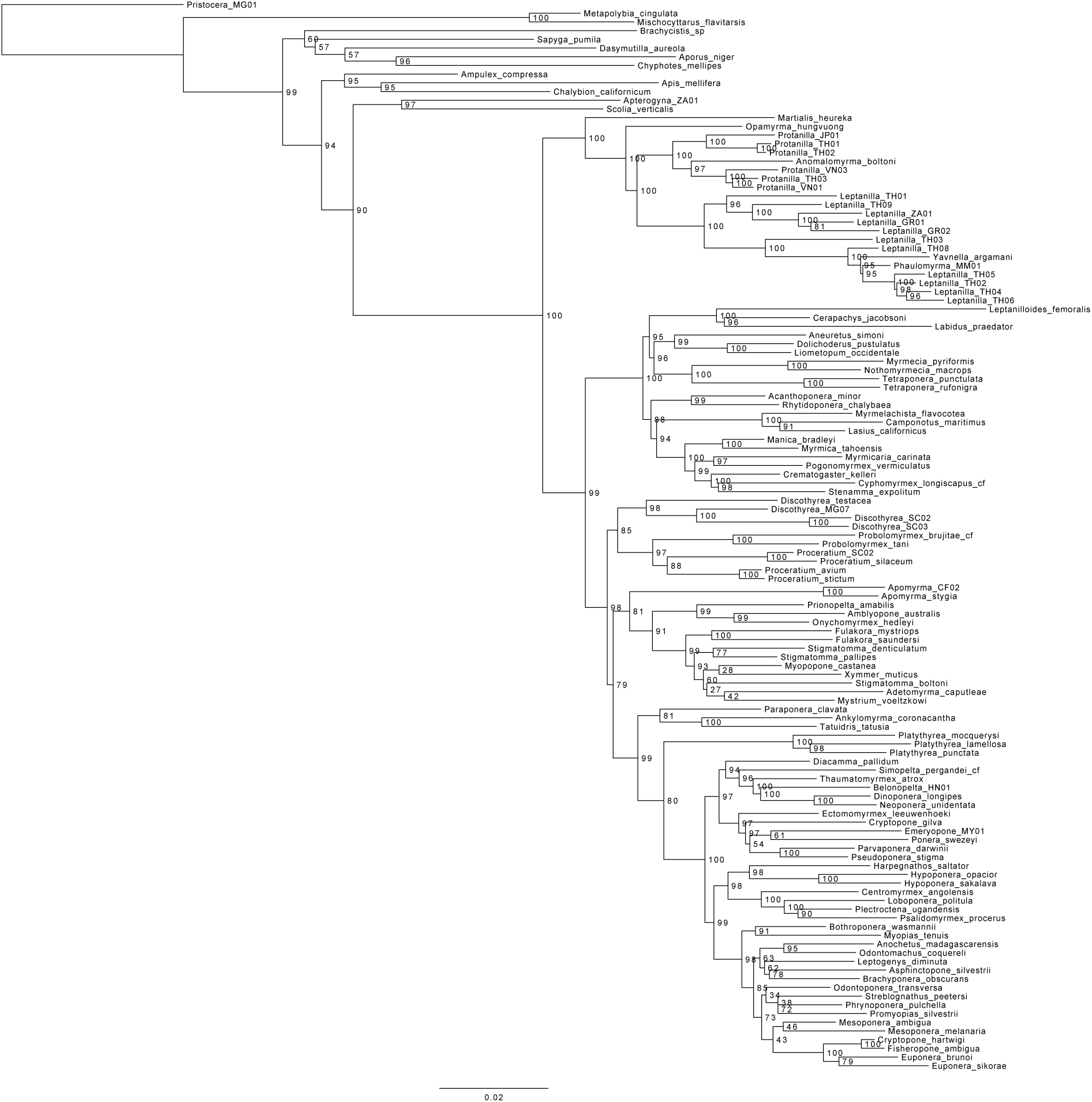
Maximum-likelihood tree inferred under greedy partitioning strategy for compositionally homogeneous datset.

**Supplementary Figure 16:**
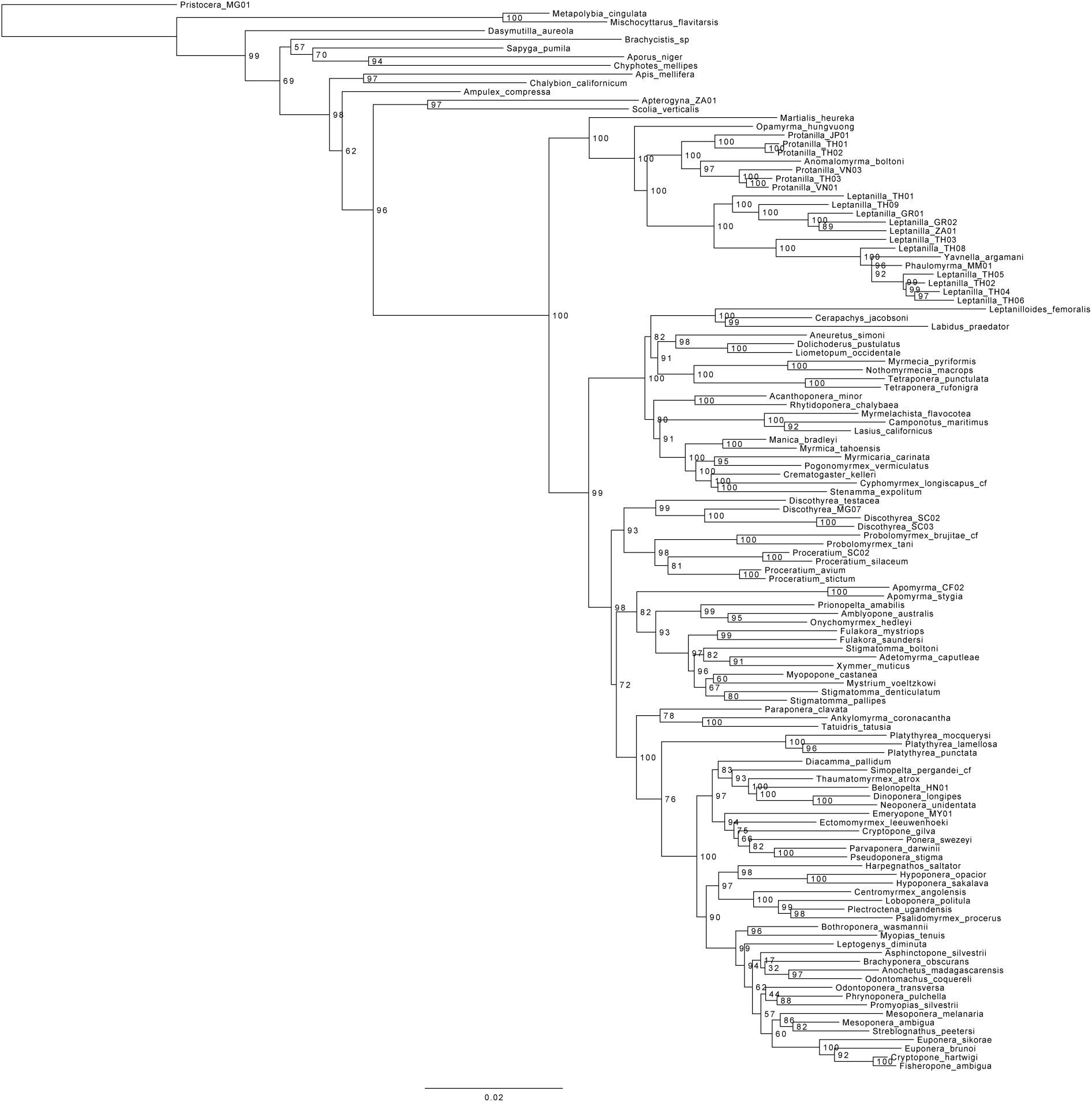
Maximum-likelihood tree inferred under k-means partitioning strategy for compositionally homogeneous datset.

**Supplementary Figure 17:**
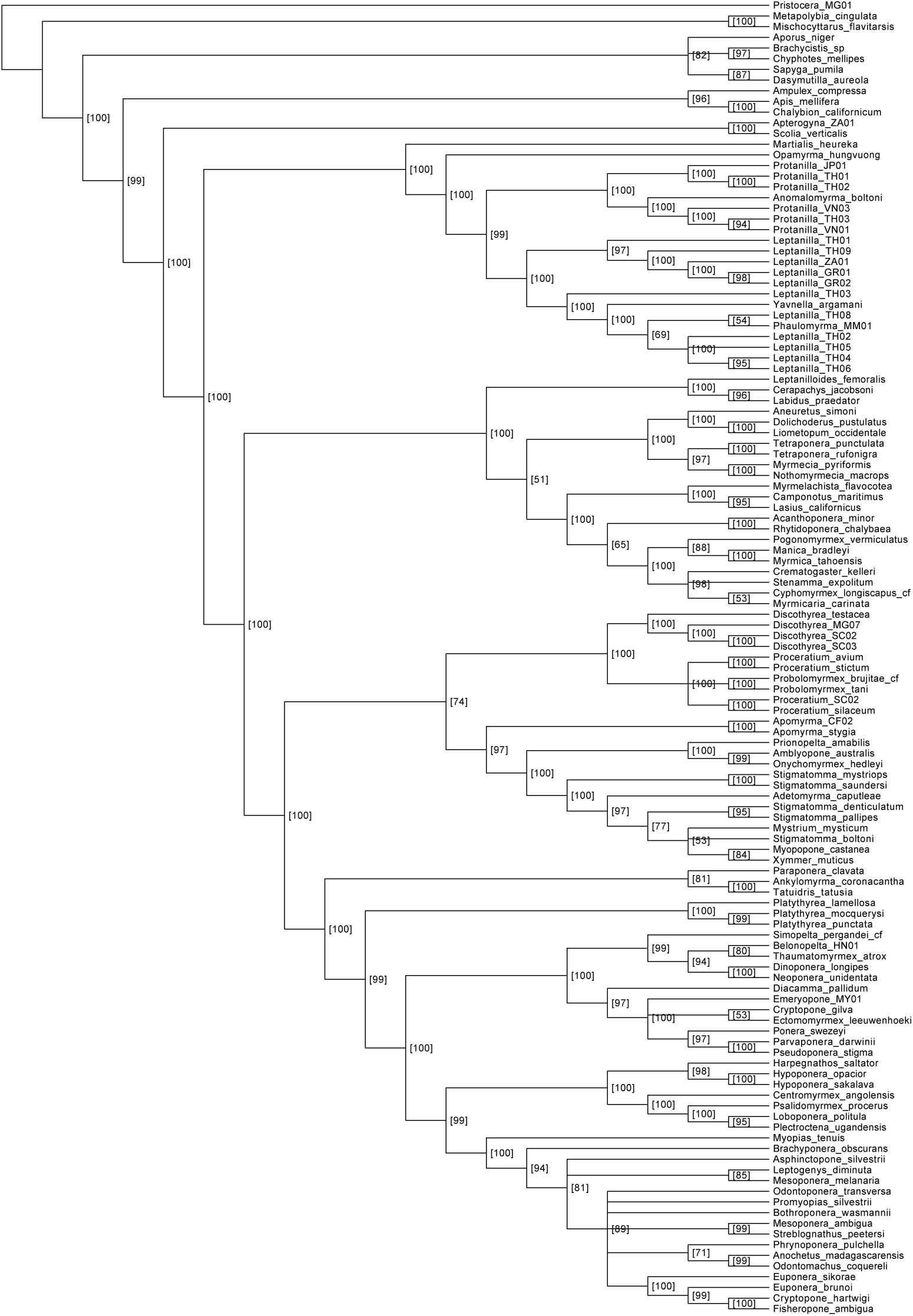
Majority-rule consensus of maximum-likelihood trees inferred from 100 simulated alignments imitating first codon positions.

**Supplementary Figure 18:**
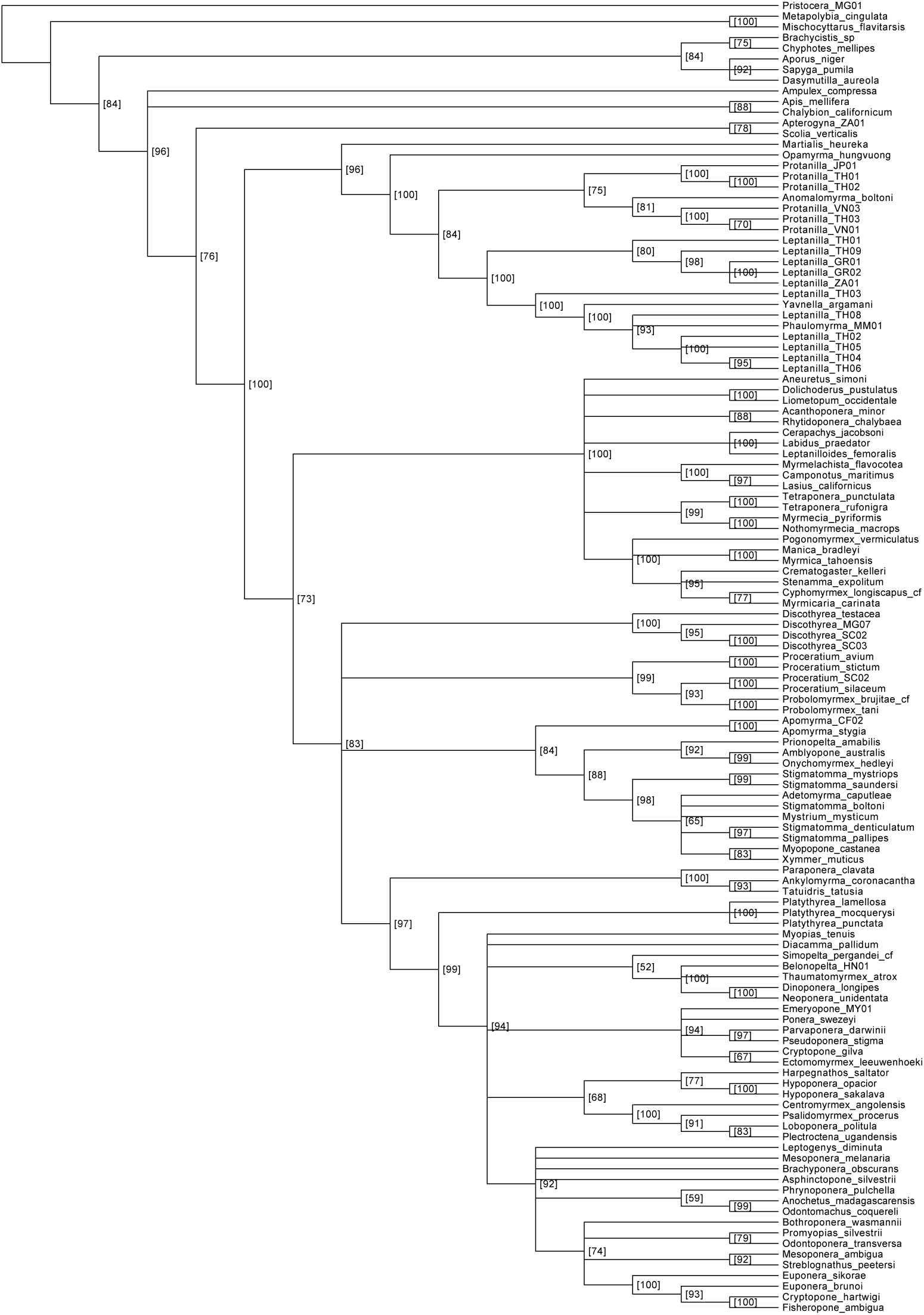
Majority-rule consensus of maximum-likelihood trees inferred from 100 simulated alignments imitating second codon positions.

**Supplementary Figure 19:**
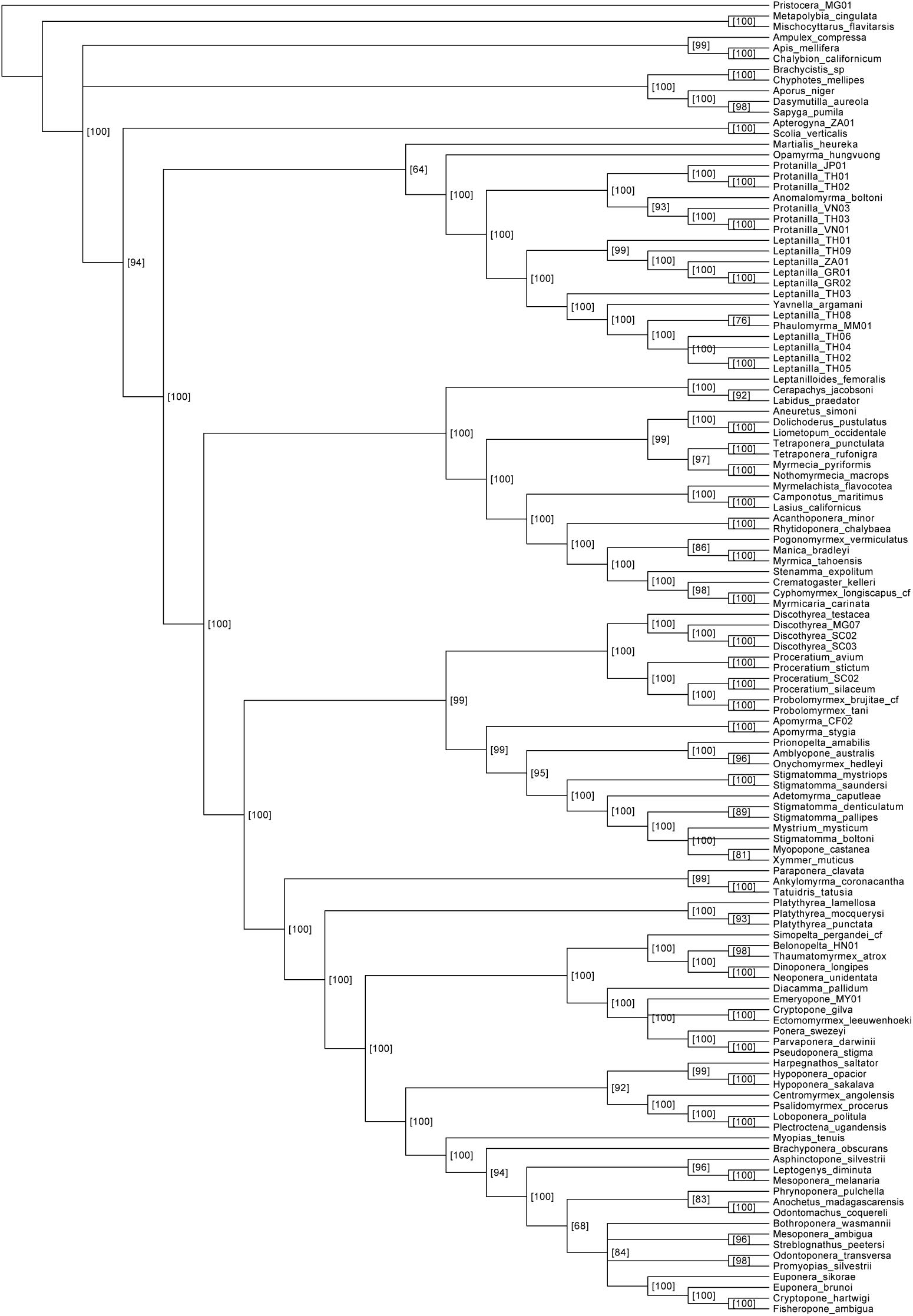
Majority-rule consensus of maximum-likelihood trees inferred from 100 simulated alignments imitating third codon positions.

**Supplementary Figure 20:**
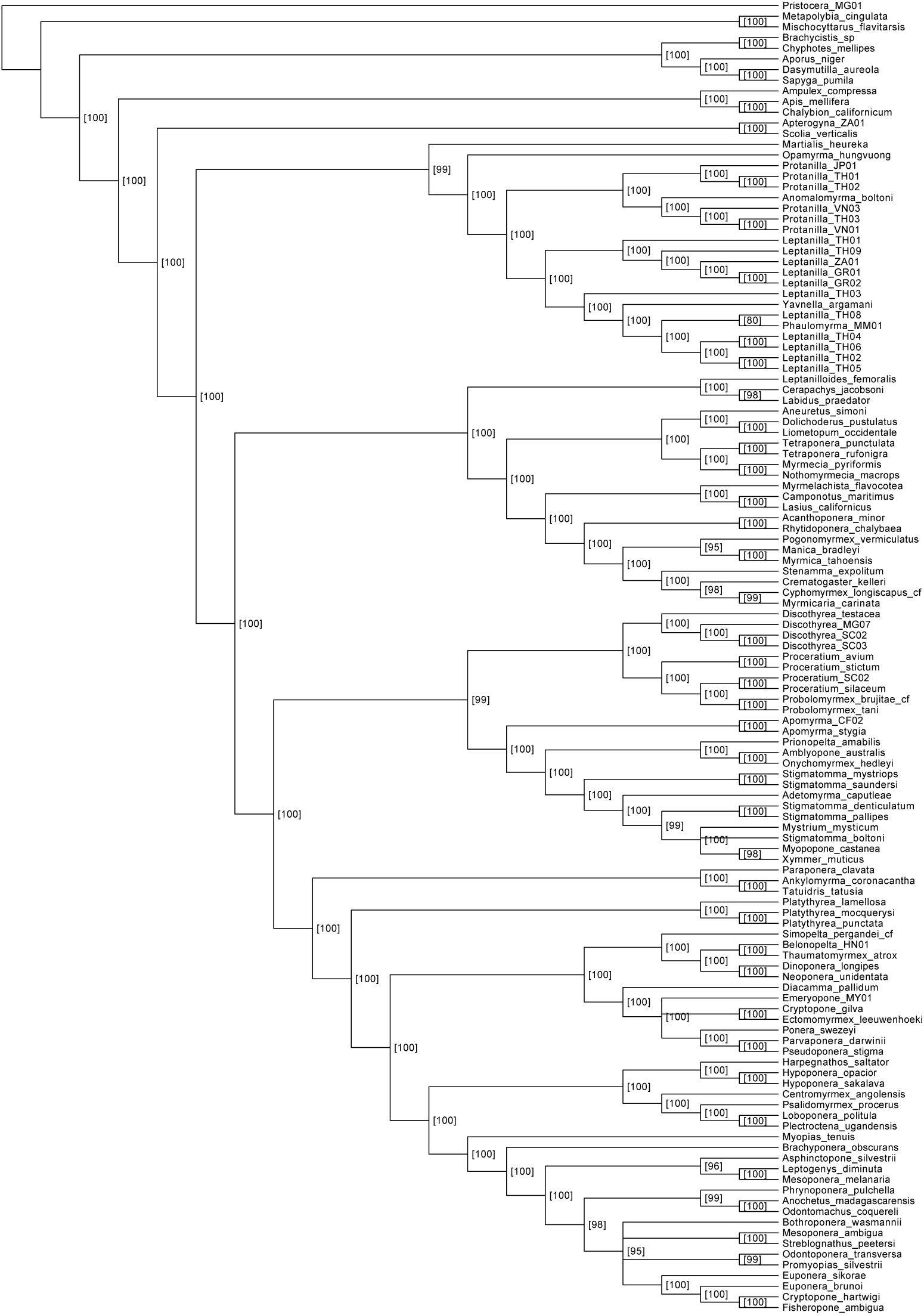
Majority-rule consensus of maximum-likelihood trees inferred from 100 simulated alignments imitating first, second, and third codon positions combined.

**Supplementary Table 1:**
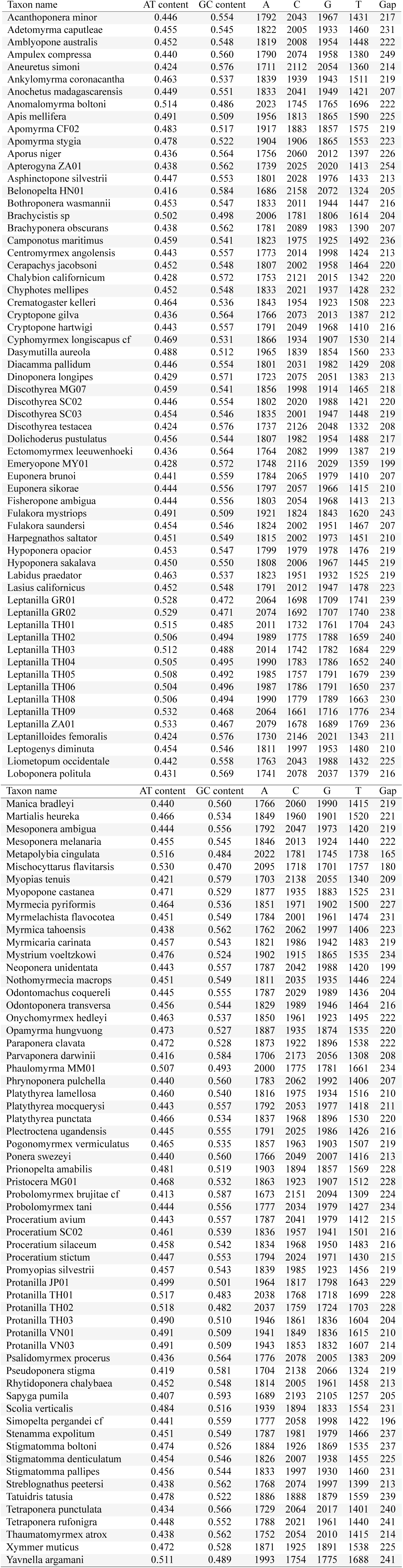
Taxon properties in the full data set. AT and GC contents are expressed as proportions and nucleotide and gap columns contain character counts. Alignment has 7,451 sites total.

**Supplementary Table 2:** Results of compositional homogeneity tests for all data partitions. P-values below 0.05 considered significant violation of homogeneity assumption.

| Partition | Corrected test p-value | Chi-squared p-value |
| --- | --- | --- |
| Full data set | 0.000 | 0.000 |
| 28S | 0.000 | 1.000 |
| abdA | 0.000 | 0.990 |
| abdA 1st pos | 0.905 | 1.000 |
| abdA 2nd pos | 0.262 | 1.000 |
| abdA 3rd pos | 0.000 | 0.000 |
| argK | 0.000 | 0.000 |
| argK 1st pos | 0.698 | 1.000 |
| argK 2nd pos | 0.987 | 1.000 |
| argK 3rd pos | 0.000 | 0.000 |
| Antp | 0.000 | 0.000 |
| Antp 1st pos | 0.847 | 1.000 |
| Antp 2nd pos | 0.520 | 1.000 |
| Antp 3rd pos | 0.000 | 0.000 |
| EF1aF2 | 0.000 | 0.000 |
| EF1aF2 1st pos | 1.000 | 1.000 |
| EF1aF2 2nd pos | 0.555 | 1.000 |
| EF1aF2 3rd pos | 0.000 | 0.000 |
| LW Rh | 0.000 | 0.000 |
| LW Rh 1st pos | 1.000 | 1.000 |
| LW Rh 2nd pos | 0.926 | 1.000 |
| LW Rh 3rd pos | 0.000 | 0.000 |
| NaK | 0.000 | 0.025 |
| NaK 1st pos | 0.558 | 1.000 |
| NaK 2nd pos | 0.460 | 1.000 |
| NaK 3rd pos | 0.000 | 0.000 |
| POLD1 | 0.000 | 0.000 |
| POLD1 1st pos | 0.004 | 1.000 |
| POLD1 2nd pos | 0.766 | 1.000 |
| POLD1 3rd pos | 0.000 | 0.000 |
| Top 1 | 0.000 | 0.000 |
| Top 1 1st pos | 0.106 | 1.000 |
| Top 1 2nd pos | 0.964 | 1.000 |
| Top 1 3rd pos | 0.000 | 0.000 |
| Ubx | 0.000 | 0.009 |
| Ubx 1st pos | 0.038 | 1.000 |
| Ubx 2nd pos | 0.202 | 1.000 |
| Ubx 3rd pos | 0.000 | 0.000 |
| Wg | 0.000 | 0.000 |
| Wg 1st pos | 0.749 | 1.000 |
| Wg 2nd pos | 0.923 | 1.000 |
| Wg 3rd pos | 0.000 | 0.000 |
| All 1st pos | 0.028 | 1.000 |
| All 2nd pos | 1.000 | 1.000 |
| All 3rd pos | 0.000 | 0.000 |

**Supplementary Table 3:**
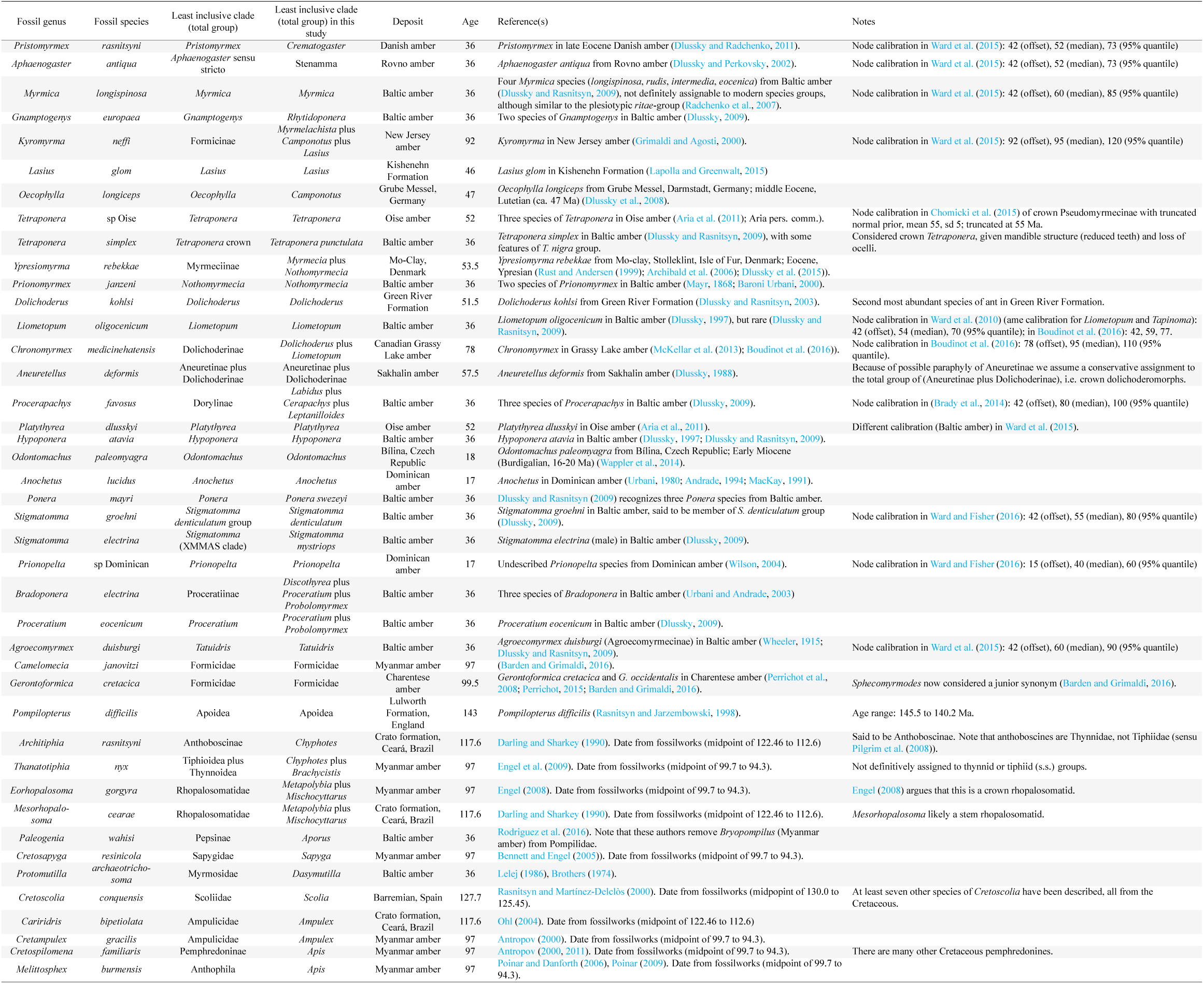
Calibration fossils used in divergence dating analysis.

**Supplementary Table 4:**
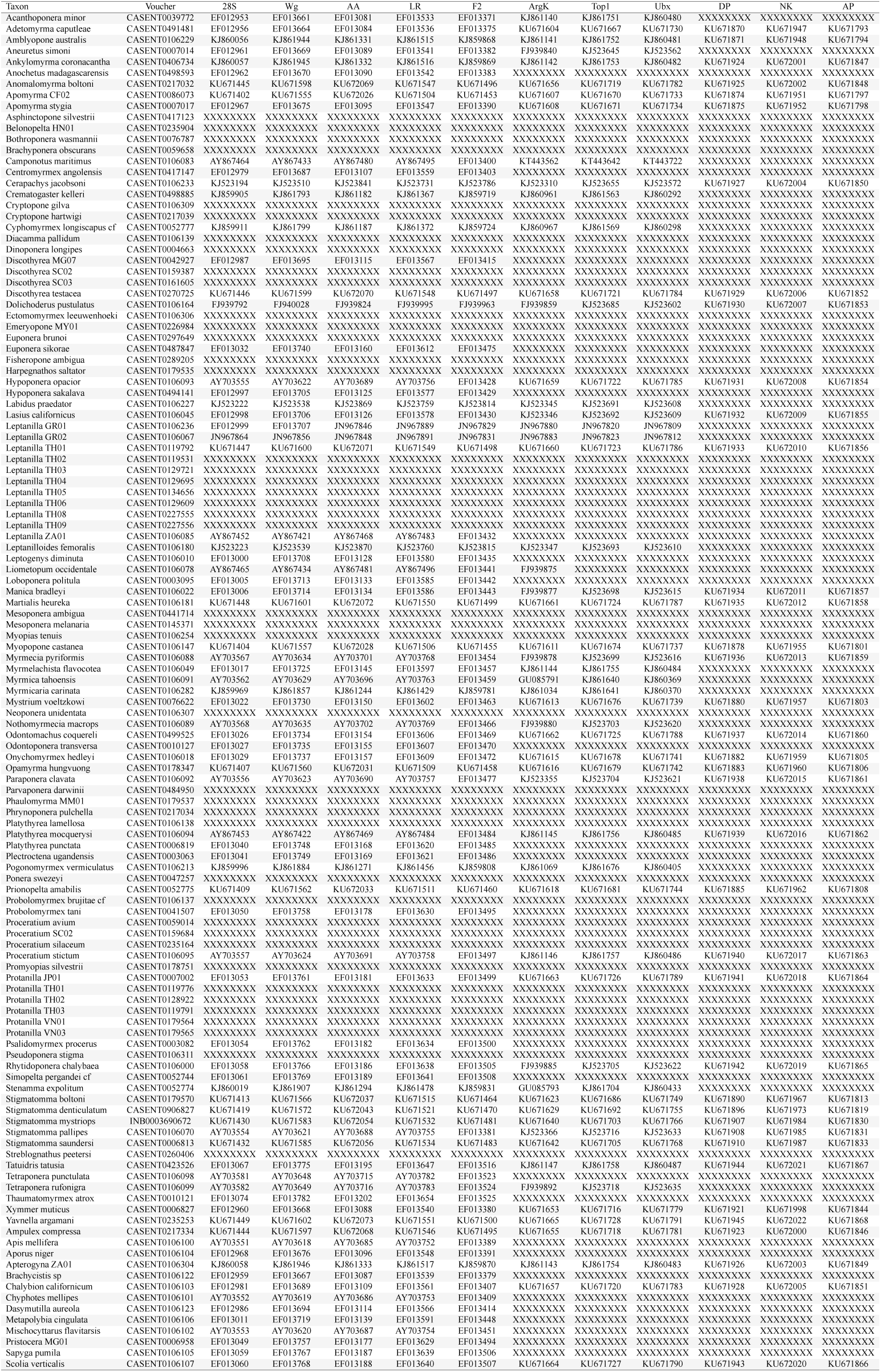
Taxa included in the study with GenBank accession numbers and specimen voucher information. Detailed collection data for each species is available by searching for the specimen code on AntWeb (www.antweb.org).

